# Transcriptional profiles of murine oligodendrocyte precursor cells across the lifespan

**DOI:** 10.1101/2024.10.27.620502

**Authors:** Dongeun Heo, Anya A. Kim, Björn Neumann, Valerie N. Doze, Yu Kang T. Xu, Yevgeniya A. Mironova, Jared Slosberg, Loyal A. Goff, Robin J. M. Franklin, Dwight E. Bergles

## Abstract

Oligodendrocyte progenitor cells (OPCs) are highly dynamic, widely distributed glial cells of the central nervous system (CNS) that are responsible for generating myelinating oligodendrocytes during development. By also generating new oligodendrocytes in the adult CNS, OPCs allow formation of new myelin sheaths in response to environmental and behavioral changes and play a crucial role in regenerating myelin following demyelination (remyelination). However, the rates of OPC proliferation and differentiation decline dramatically with aging, which may impair homeostasis, remyelination, and adaptive myelination during learning. To determine how aging influences OPCs, we generated a novel transgenic mouse line that expresses membrane-anchored EGFP under the endogenous promoter/enhancer of Matrilin-4 (*Matn4-mEGFP*) and performed high-throughput single-cell RNA sequencing, providing enhanced resolution of transcriptional changes during key transitions from quiescence to proliferation and differentiation across the lifespan. Comparative analysis of OPCs isolated from mice aged 30 to 720 days, revealed that aging induces distinct inflammatory transcriptomic changes in OPCs in different states, including enhanced activation of HIF-1α and Wnt pathways. Inhibition of these pathways in acutely isolated OPCs from aged animals restored their ability to differentiate, suggesting that this enhanced signaling may contribute to the decreased regenerative potential of OPCs with aging. This *Matn4-mEGFP* mouse line and single-cell mRNA datasets of cortical OPCs across ages help to define the molecular changes guiding their behavior in various physiological and pathological contexts.

---

Aging is accompanied by a progressive decline in the functional capabilities and restorative capacity of the brain, resulting in increased susceptibility to neurodegenerative disease. Cellular dysregulation within brain circuits is normally mitigated by glial cells; however, glia are also vulnerable to metabolic stress, somatic mutations, and cellular senescence that increase with aging^1^. Phenotypic changes in glia can disrupt ion and neurotransmitter homeostasis, increase inflammation, and induce the release of toxic factors that disrupt function and impair the survival of surrounding neurons^2–4^. Understanding how different glial cells are influenced by the aging brain environment and contribute to this declining brain resilience requires longitudinal assessments of their molecular characteristics across the lifespan.

Although the brain has a limited capacity to regenerate neurons damaged through trauma or disease, it retains a population of lineage-restricted progenitors that have the capacity to develop into myelin-forming oligodendrocytes. These oligodendrocyte precursor cells (OPCs) continually produce new oligodendrocytes in the adult CNS, increasing total myelin content within brain circuits and altering the pattern of myelin along distinct neuron subtypes^5–7^. OPC differentiation can be enhanced through motor training and enhanced sensory experience^8,9^, an adaptive form of myelination that may contribute to functional changes in neural circuits necessary for learning. OPCs also play a critical role in regenerating oligodendrocytes destroyed by trauma, stroke, and diseases such as multiple sclerosis (MS). However, the ability to form new oligodendrocytes declines with age^10,11^, which may contribute to remyelination impairment in MS and myelin loss in aging-associated dementia. In addition, recent studies indicate that some oligodendrocyte lineage cells undergo aging-associated cellular senescence^12^ and upregulate antigen presentation pathways in neurodegenerative diseases, such as Alzheimer’s disease (AD) and MS^13–15^. However, the molecular mechanisms underlying these aging-associated changes in their dynamics and lineage progression remain poorly understood.

Despite their persistence and impact on regenerative processes, OPCs constitute only a small proportion of brain cells (approximately 2-3% in gray matter, 5% in white matter)^16^. Moreover, oligodendroglia exist in a developmental continuum from cycling progenitors (OPCs) to terminally differentiated cells (oligodendrocytes), complicating the assessment of transcriptional changes within cells at each stage when assessed from small samples. To increase the resolution of state-dependent transcriptional changes and aging-associated alterations in OPCs, we generated a novel line of transgenic knock-in mice (*Matn4-mEGFP*), in which OPCs throughout the CNS express membrane-anchored EGFP and used these animals to isolate and perform single-cell mRNA sequencing (scRNA-seq) of OPCs from the cerebral cortex of young, adult, and aged mice. Enrichment of OPCs in these samples provided greater resolution of transcriptional changes associated with their proliferation and differentiation, and revealed distinct features associated with aging, such as enhancement of HIF-1α and Wnt signaling, and upregulation of complement expression. By performing *in vitro* pharmacological manipulations of OPCs isolated from aged mice, we show that the inhibition of HIF-1α and Wnt signaling pathways markedly enhances their ability to differentiate. Together, these new transgenic mice and transcriptional information of cortical OPCs across the lifespan provide a means to identify new strategies to restore the regenerative potential of these progenitors in aging and diverse neurodegenerative diseases.

## Results

### *Matn4*-*mEGFP* mice enable *in vivo* visualization and selective isolation of OPCs from the CNS

It has been difficult to assess the diversity of OPCs or the molecular changes they exhibit as the brain ages using bulk or unbiased isolation approaches, due to their relatively low abundance. In addition, many molecular markers used to identify OPCs, such as NG2, PDGFRα, and Olig2, are not specific to OPCs. To develop an enrichment strategy for OPCs, we searched available transcriptomic datasets to identify genes selectively expressed by these cells. We determined that mRNA encoding *Matn4,* an extracellular matrix protein that regulates stress-induced proliferation of hematopoietic stem cells^17^, is highly enriched in OPCs compared to other neural cells in the cerebral cortex^18^ (Extended Data Fig. 1a). Using CRISPR-Cas9 gene editing, we generated a novel transgenic mouse line by inserting a membrane-anchored EGFP (mEGFP) sequence at the first coding exon of *Matn4* (Fig. 1a; Extended Data Fig. 1b; Supplementary Data 1). In these *Matn4-mEGFP* (*Matn4^mEGFP/+^*) mice, small cells with radially oriented processes expressed EGFP and were organized in a grid-like pattern with non-overlapping territories, consistent with the known morphology and distribution of OPCs in the CNS^19^ (Fig. 1b). Importantly, perivascular cells (pericytes and perivascular fibroblasts), which also express NG2 and PDGFRα^20,21^, were not EGFP immunoreactive (EGFP+) in these animals (Fig. 1c), demonstrating the utility of this mouse line for unambiguous OPC identification. Immunocytochemical analysis in the brain, spinal cord, and optic nerve of heterozygous *Matn4-mEGFP* mice revealed that these cells also expressed NG2, a proteoglycan expressed by OPCs (Fig. 1d; Extended Data Fig. 1c) and that this specificity (% of NG2+/all EGFP+ cells) and efficiency (% of EGFP+/NG2+ PDGFRα+ OPCs) of OPC labeling were preserved in aged (P720) *Matn4-mEGFP* mice (Fig. 1e-g). Further analysis revealed that apart from OPCs, mEGFP was also expressed by hippocampal granule cells^22^ and by neurons in the somatosensory barrel field and retrosplenial cortex (Extended Data Fig. 1d), consistent with previous transcriptomic studies^22,23^. EGFP signal was not detected in Iba1+ microglia or GFAP+ astrocytes in the brain (Extended Data Fig. 1e).

**Figure 1.**
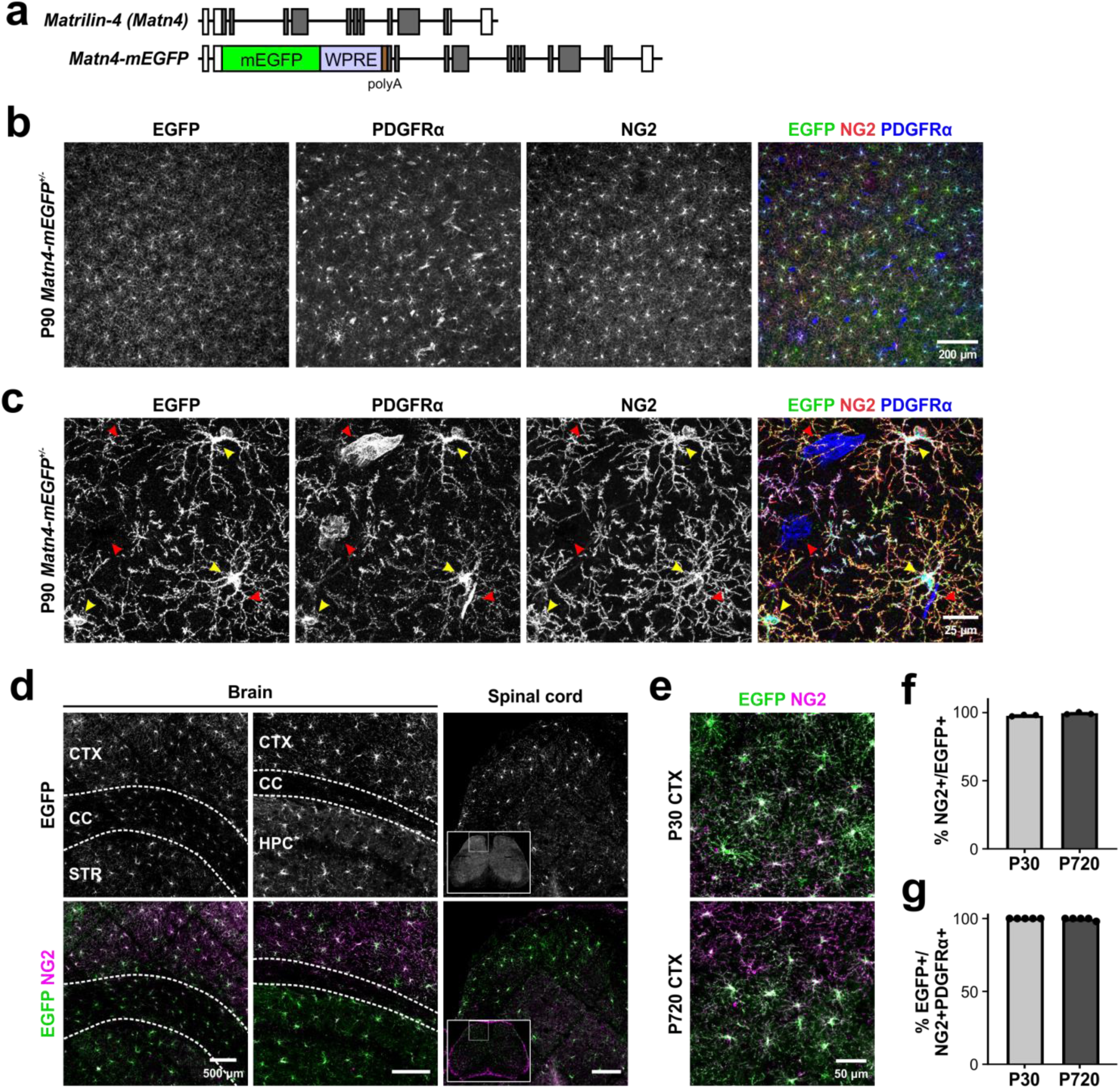
Generation of an oligodendrocyte precursor cell (OPC) reporter mouse line: *Matn4-mEGFP*. **a.** Schematics of *Matn4-mEGFP* mouse line where membrane anchored EGFP (mEGFP), WPRE, and polyA sequences were knocked into the first coding exon of *Matn4.* **b.** Confocal images of flattened *Matn4-mEGFP* mouse cortex co-immunostained for NG2 and PDGFRα. **c.** Higher magnification confocal images show the specificity of *EGFP* expression by NG2+ PDGFRα+ OPCs (yellow arrowheads), but not PDGFRα+ perivascular fibrocytes (red arrowheads). **d.** EGFP+ OPCs in *Matn4-mEGFP* mice also express NG2 in the corpus callosum (CC), striatum (STR), hippocampus (HPC), and spinal cord. **e.** The specificity of labeling OPCs does not change with aging in the cortex (CTX). **f.** Quantification of labeling specificity in *Matn4-mEGFP* mice, illustrating the percentage of EGFP+ cells that are also NG2+ in the cortex at P30 and P720. **g.** Quantification of labeling efficiency, illustrating the percentage of NG2+ PDGFRα+ OPCs expressing EGFP in the cortex at P30 and P720.

To determine if these mice could also be used to study the dynamics of OPCs *in vivo*, we implanted cranial windows over the primary motor cortex (M1) of *Matn4-mEGFP* mice and performed two-photon, time-lapse fluorescence imaging. Consistent with the histological analysis, in young adult *Matn4-mEGFP* mice, OPCs were visible throughout the upper 200 μm of area M1 and exhibited dynamic behavior, consisting of filopodial extension and retraction, process reorientation, soma translocation, and cell division, comparable to that described previously in *NG2-mEGFP* mice^24^ (Extended Data Fig. 1f, Supplementary Videos 1 and 2). Together, these results indicate that OPCs throughout the CNS express mEGFP in *Matn4-mEGFP* mice, providing a means to visualize and isolate these cells from the intact CNS to examine their phenotypic changes during aging.

### OPCs in the cerebral cortex exist in transcriptionally distinct states

To determine how aging influences gene expression by OPCs, we used fluorescence-activated cell sorting (FACS) to isolate OPCs from the cerebral cortex of *Matn4-mEGFP* mice using endogenous EGFP fluorescence at four different ages, spanning young adult (P30), adult (P180), middle age (P360), and aged (P720) stages of life. Single-cell droplets were generated from dissociated whole cortices using the 10x Chromium controller (10x Genomics) and libraries were prepared using the 3’ gene expression platform. Uniquely barcoded libraries were pooled and sequenced to an approximate depth of 50,000 reads/cell (5 batches, 20 samples) (Fig. 2a). More than 98% of all cells in these samples were oligodendrocyte lineage cells, based on the expression of genes associated with OPCs and oligodendroglia (*Cspg4, Pdgfra, Olig2, Enpp6*) (Extended Data Fig. 2a) and the lack of mRNA transcripts associated with other cell types, such as *Slc17a7* (excitatory neurons)*, Gad2* (inhibitory neurons)*, Aldh1l1* (astrocytes)*, Cx3cr1* (microglia)*, Cldn5* (endothelial cells), and *Vtn* (pericytes) (Extended Data Fig. 2b). After removing cells with >5% of mitochondrial gene read ratio from the dataset (Extended Data Fig. 2c), this sampling provided high-quality transcriptional profiles from 38,807 OPCs (Fig. 2b), a 17-fold increase from previous aging studies obtained through unbiased, bulk sampling of all neural cells in the brain^25^.

**Figure 2.**
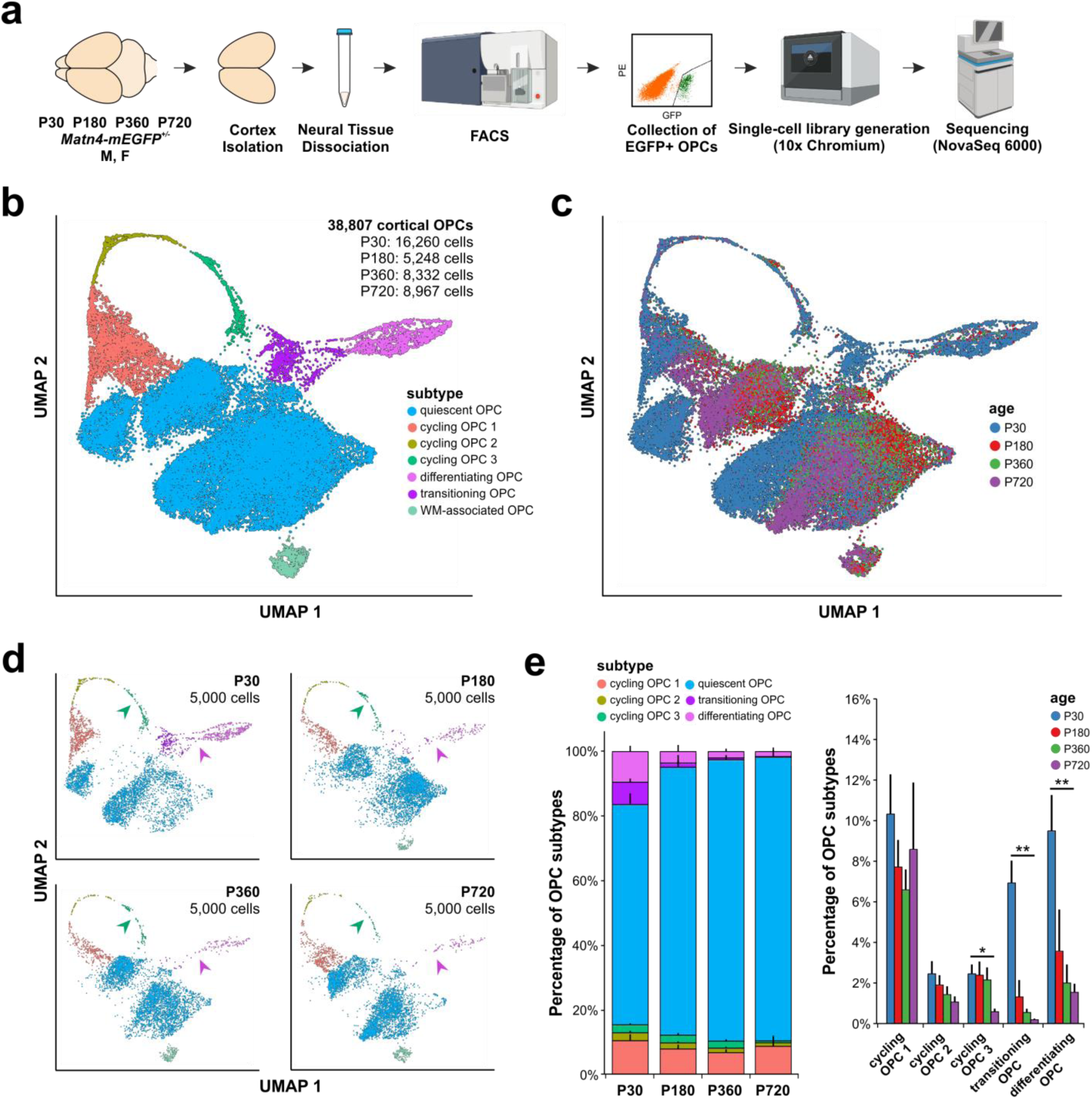
Single cell RNA-seq analysis of mouse cortical OPCs across the lifespan. **a.** Workflow for generating the 10x Chromium single cell RNA-seq dataset from P30, P180, P360, and P720 mouse cortical OPCs acutely isolated from *Matn4-mEGFP* mice (illustrations created in BioRender: https://BioRender.com/z82n750). **b.** UMAP plot of 38,807 mouse cortical OPCs from four timepoints, colorized by their identified subtypes (*Quiescent OPC*, *Cycling OPC 1*, *Cycling OPC 2*, *Cycling OPC 3*, *Differentiating OPC*, *Transitioning OPC*, and *WM-associated OPC*). **c.** UMAP plot of the dataset colorized by their four age groups (blue: P30, red: P180, green: P360, and purple: P720). **d.** Separate UMAP plots for different age groups colorized by their subtypes. 5,000 cells from each age group were randomly selected and plotted. The arrowheads denote cycling OPC 3 (green) and transitioning/differentiating OPC (pink) subtypes. **e.** Proportions of each OPC subtype across four age groups show that the quiescent OPC subtype increases in proportion due to a statistically significant reduction in the proportions of cycling OPC 2, cycling OPC 3, transitioning OPC, and differentiating OPC subtypes with aging (simple linear regression, n=6, 4, 5, 5, * p-value < 0.05, ** p-value < 0.01).

With the increased resolution provided by this extensive sampling of OPCs, we were able to identify OPCs in distinct states, including a transient population of cells that were in the initial transition from progenitor to premyelinating oligodendrocyte. After initial unbiased clustering, we grouped OPCs into four different populations (*Cycling*, *Differentiating*, *Transitioning*, and *Quiescent*) based on the differential expression of genes previously associated with cell division (*Top2a, Mcm3, Mki67*) and actively differentiating OPCs (*Bcas1*^26^*, Enpp6*^27^*, 9630013A20Rik/*LncOL1^28^) (Fig. 2b; Extended Data Fig. 2d; Supplementary Data 2). Fluorescent *in situ* hybridization against *Top2a* and *LncOL1* in young (P9) and adult (P74) mouse brains showed that these transient OPC states can be visualized using these marker genes (Extended Data Fig. 3). The *Transitioning OPC* subtype (highly expressing *Gap43, Rplp0*) (Extended Data Fig. 2d; Supplementary Data 2) was identified and annotated after a subsequent analysis of the subset of *Quiescent* OPCs that immediately precede the *Differentiating OPC* subtype. We also identified a small population of OPCs (*WM- associated*) (Fig. 2b) that was enriched for genes previously associated with OPCs in white matter (e.g. *Ednrb*^29^), suggesting that they may reflect the inclusion of a portion of the corpus callosum during dissection or a population of OPCs in the deeper cortical layers that become heavily myelinated over the course of aging. This *WM-associated OPC* subtype also highly expressed *Clusterin* (*Clu*), which has previously been suggested to underlie OPC heterogeneity in the adult mouse brain^30^.

When pooled across ages, the *Cycling OPC* subtype represented 13.1% of all OPCs (P30: 15.2 ± 2.3%, P180: 12.0 ± 1.6%, P360: 10.2 ± 1.8%, P720: 10.2 ± 3.3%) and the *Transitioning* and *Differentiating* subtypes comprised 7.2% of all cells (P30: 16.4 ± 2.7%, P180: 4.9 ± 2.9%, P360: 2.5 ± 0.8%, P720: 1.7 ± 0.4%). The remaining cells, which represented the majority of OPCs at all ages (P30: 68.3 ± 3.5%, P180: 83.1 ± 3.8%, P360: 87.3 ± 1.4%, P720: 88.1 ± 3.0%), were termed *Quiescent*, as they lacked gene signatures associated with these dynamic behaviors. Plotting a random selection of 5,000 OPCs from each age highlights the decline in *Cycling*, *Transitioning*, and *Differentiating* OPCs with aging (Fig. 2d). This progressive decrease in the proportion of cycling and differentiating cells with age (Fig. 2e) is consistent with previous *in vivo* assessments using BrdU/EdU incorporation and genetic fate tracing^31,32^, as well as bulk RNA-seq of OPCs from mouse brain^33^.

To explore the abundance of these different OPC subtypes in the human brain, we analyzed a human single-nucleus RNA-seq (snRNA-seq) dataset that includes a large population of human OPCs across aging^34^ (Extended Data Fig. 4a). In these samples, small populations of *Cycling* and *Differentiating* OPC subtypes, in addition to *Quiescent OPC* subtypes were present (Extended Data Fig. 4b). We were able to demonstrate that different human OPC subtypes expressed known marker genes, as well as those marker genes identified in our mouse dataset (Extended Data Fig. 4c). We then used scCoGAPS^35^ and projectR^36^ to project OPC subtype-specific gene patterns identified in our mouse OPC scRNA-seq dataset onto the human OPC snRNA-seq dataset. Mouse gene patterns associated with either the *Cycling OPC* (pattern 5) or *Differentiating OPC* (pattern 2) subtype could successfully be projected onto the human dataset to unbiasedly reveal corresponding *Cycling* and *Differentiating OPC* subtypes (Extended Data Fig. 4d-e; Supplementary Data 6). This analysis suggests that the global transcriptional changes that occur during the transitions from *Quiescent OPC* to *Cycling* and to *Differentiating OPC* subtypes may be evolutionarily conserved between mouse and human. Further, this comparison highlights the value of having a robust, comprehensive transcriptomic dataset of OPCs for resolving transient cell stages in other samples. The complete OPC transcriptomic dataset generated from *Matn4-mEGFP* mice can accessed through the cellxgene web interface^37^ (https://tinyurl.com/aging-opcs), allowing visualization of gene expression by distinct subtypes of OPCs at these different ages.

### OPCs in older mice exhibit reduced progression through the cell cycle

OPCs exhibit robust homeostasis in the adult CNS, rapidly proliferating in response to the loss of these cells through differentiation or death to maintain their density^24^. To define the transcriptional changes exhibited by OPCs at different points in the cell cycle, we recursively analyzed cycling OPCs to identify subclusters based on the expression of stage-specific genes, providing enhanced resolution of transcriptional changes that occur in these cells during division (Fig. 3a). Cycling OPCs were subdivided into three discrete groups using the Tricycle (Transferable Representation and Inference of Cell Cycle) R/Bioconductor software, which computationally predicts cell cycle positions using the known dynamics of different cell cycle-associated genes^38^: G1/G0-phase (*Cycling OPC 1*; 73.4%), G2/M-phase (*Cycling OPC 2*; 12.3%), and M-phase (*Cycling OPC 3*; 14.2%) (Fig. 3b). Tricycle analysis confirmed a prominent reduction of OPCs in G2/M- and M-phases (*Cycling OPC 2* and *Cycling OPC 3*) in the aged (P720) compared to the young (P30) brain (Fig. 3c). This age-associated reduction in proliferating OPCs specifically in G2/M-phase corroborates previous observations using fluorescence indicators^29^. Consistent with the tricycle prediction, pseudotime analysis identified progression of OPCs from *Cycling OPC 1* to *3* during division (Fig. 3d). By leveraging the high number of cells undergoing proliferation, we were able to establish a high-resolution pseudotemporal trajectory to identify different cell cycle-associated genes as OPCs progress through cell division (Fig. 3e). Despite the significant decrease in the proportion of *Cycling OPC 2* and *3* subtypes in the older mouse cortex, OPCs that reached these stages had remarkably similar transcriptomic profiles (Extended Data Fig. 5a), suggesting that the age-dependent decline in OPC proliferation reflects changes in prior states, and that once committed, they follow a consistent transcription program for DNA synthesis and mitosis.

**Figure 3.**
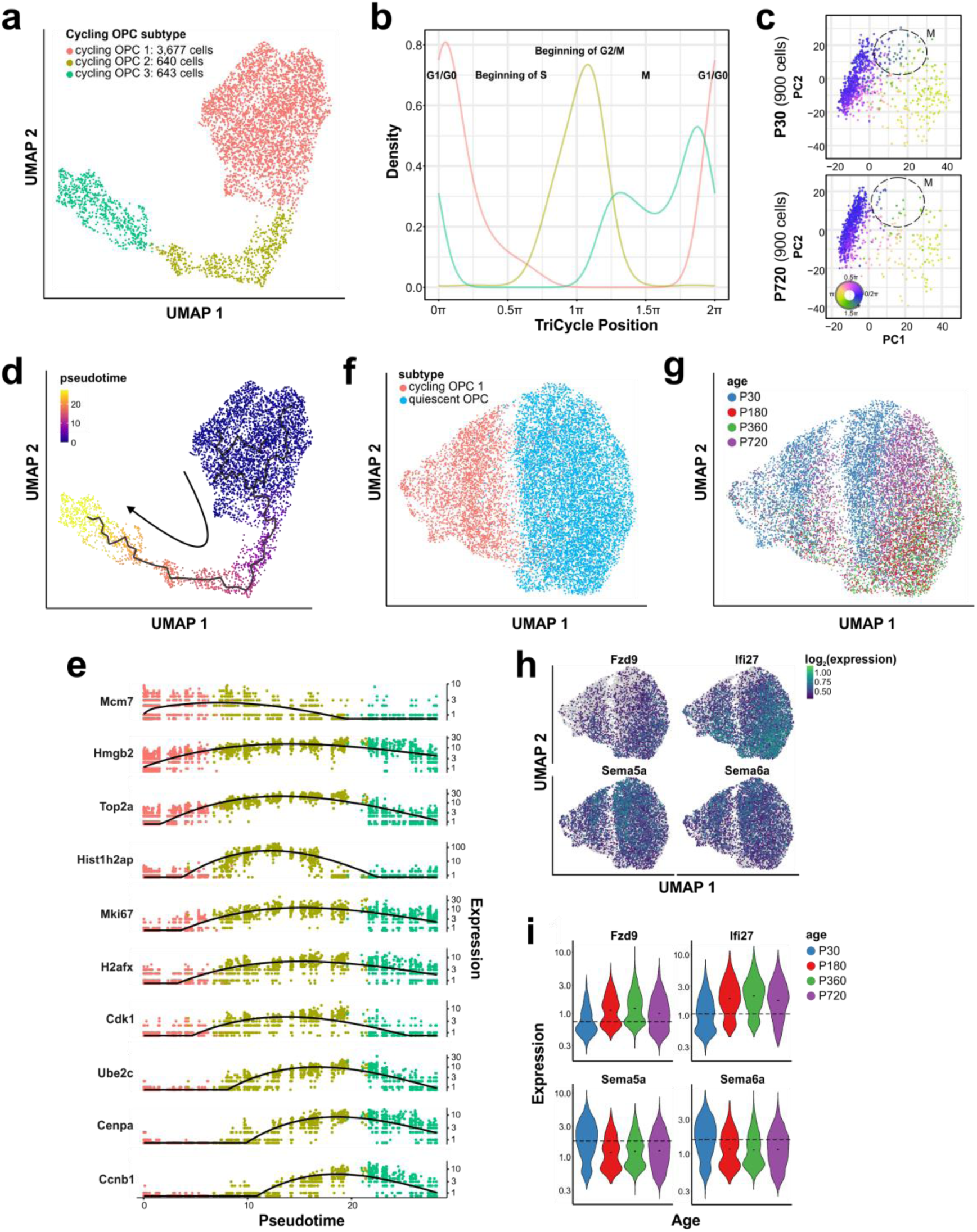
Different cycling OPC subtypes represent different stages of the cell cycle. **a.** UMAP plot of cycling OPCs (n=4,960) colorized by three different cycling OPC subtypes. **b.** TriCycle analysis shows that different cycling OPC subtypes correspond to different cell cycle stages (G1/G0: 0π/2π, S: 0.5π, G2/M: 1π, M: 1.5π). **c.** P720, aged group shows a dramatic loss of those cycling OPCs that are predicted to be undergoing mitosis (M-phase) (manually encircled for visualization). **d.** UMAP plot of cycling OPCs colorized by pseudotime originating from *Cycling OPC 1* (G1/G0-phase). **e.** Expression of different cell-cycle related genes in cycling OPCs across the pseudotime, colorized by their subtypes (*Cycling OPC 1*, *Cycling OPC 2*, and *Cycling OPC 3*). **f.** UMAP plot of *Cycling OPC 1* (G1/G0-phase) and immediately anteceding quiescent OPCs colorized by their subtypes. **g.** f colorized by different age groups (P30, P180, P360, and P720). **h.** UMAP plots of the top four genes (*Fzd9, Ifi27, Sema5a,* and *Sema6a*) differentially expressed across aging in quiescent OPCs that directly antecede cycling OPCs. **i.** Expression violin plots of the top four genes in h across the four different age groups. *Fzd9* and *Ifi27* are statistically significantly upregulated with aging whereas *Sema5a* and *Sema6a* are significantly downregulated (monocle3 linear regression, q-value < 0.001).

To determine what differences account for these aging-dependent changes in cell cycle entry, we performed a comparative transcriptional analysis of cells in *Cycling OPC 1* and *Quiescent* stages immediately preceding the *Cycling OPC 1* stage (Fig. 3f,g). Although it is bioinformatically difficult to differentiate between G0- and G1-phases of the cell cycle^38^, the *Cycling OPC 1* subtype expresses canonical cell cycle-associated genes, such as *Hells* and *Mcm2-7* (Supplementary Data 2) whereas *Quiescent* OPCs do not. Using non-negative matrix factorization (NMF) to identify transcriptional features associated with distinct biological processes^39^, we found aging-associated patterns of co-regulated genes that may explain the separation of older OPCs from young, P30 OPCs (Extended Data Fig. 5b). Among the highly weighted genes from the aged gene pattern (Pattern_2) (Extended Data Fig. 4b), *Ifi27* and *Fzd9* were statistically significantly upregulated in the P180-720 cells in *Cycling OPC 1* and *Quiescent* OPC subtypes that were poised to enter the G1-phase of the cell cycle (Monocle likelihood ratio test, 0.05% FDR) (Fig. 3h,i). *Ifi27*, which encodes for Interferon Alpha Inducible Protein 27, has been primarily characterized in peripheral tissues as one of the downstream targets of type-I interferon (IFN-I) signaling. Notably, IFN-I signaling is upregulated in aged human and mouse choroid plexus, which may critically link peripheral immunity with brain senescence^40^. *Fzd9* (Frizzled Class Receptor 9) has been shown to be regulated by m6A RNA methylation in oligodendrocytes^41^ and detected in chronic active MS lesions, where it has been proposed as a negative regulator of OPC differentiation^42^.

Expression of the semaphorins *Sema5a* and *Sema6a* were significantly downregulated in aged (P180- 720), compared to young (P30) cells in *Cycling OPC 1* and *Quiescent* OPC subtypes (Monocle likelihood ratio test, 0.05% FDR) (Fig. 3h,i). Semaphorins comprise a large family of secreted guidance molecules that influence neuronal morphogenesis and control OPC migration^43^. Both *Sema5a* and *Sema6a* are expressed by oligodendroglia^44^ and *Sema6a* promotes OPC differentiation during development^45^, raising the possibility that a reduction of this signaling may bias OPCs away from lineage progression toward quiescence and proliferation. Oligodendroglial *Sema5a* has been shown to inhibit the outgrowth and regeneration of retinal ganglion cell axons (RGCs)^46^, but its role in controlling the cell-intrinsic behavior of OPCs has not yet been explored. Together, this comparative transcriptional analysis reveals possible contributors to the aging-dependent decline in OPC proliferation.

### Aging-associated transcriptional changes associated with oligodendrogenesis

OPCs continue to differentiate in the adult CNS, providing new oligodendrocytes that increase myelin content along axons^5,6,9^. However, both the generation of new oligodendrocytes and the ability to regenerate oligodendrocytes after injury or disease decline with age^11,12,31,47^. This behavior was evident in OPCs isolated from the cerebral cortex of *Matn4-mEGFP* mice, as a reduction of OPCs within the *Transitioning* and *Differentiating* stages between P30 and the older mice (Fig. 2d,e, *purple* and *magenta*). Although there was a progressive decline in cells in these states from P180 to P720 (Fig. 2e), the proportion of cells in the *Differentiating* group was not significantly different across these older time points, although it trended lower with increasing age (P180: 2.3 ± 2.0%, P360: 2.0 ± 0.9%, P720: 1.4 ± 0.4%, simple linear regression). This persistence highlights that some OPCs retain the ability to execute transcriptional changes associated with oligodendrogenesis over the adult lifespan where this process seems to progress at a constant, albeit low rate. To define transcriptional changes associated with this crucial state transition at a higher cellular resolution, we computationally isolated and performed re-clustering of *Quiescent OPCs* immediately preceding the *Transitioning* and *Differentiating OPC*s (Fig. 4a-c). Using NMF, we identified 14 different patterns of co-regulated gene expression (Fig. 4d). *Transitioning OPC*s were defined by the transcriptomic signature of module 13, which showed a diverse set of ribosomal protein (*Rpl* and *Rps*) genes to have high gene weights, consistent with the dramatic post-transcriptional changes that quiescent OPCs require to undergo differentiation (Fig. 4e; Supplementary Data 3). In addition, OPCs in later stages of differentiation were defined by module 2, with many zinc finger proteins (e.g. *Zfp36*, *Zfp365*) and canonical differentiation-associated genes, such as *Myrf* and *Bcas1*^26,48^ demonstrating high weights in this module (Fig. 4e; Supplementary Data 3).

**Figure 4.**
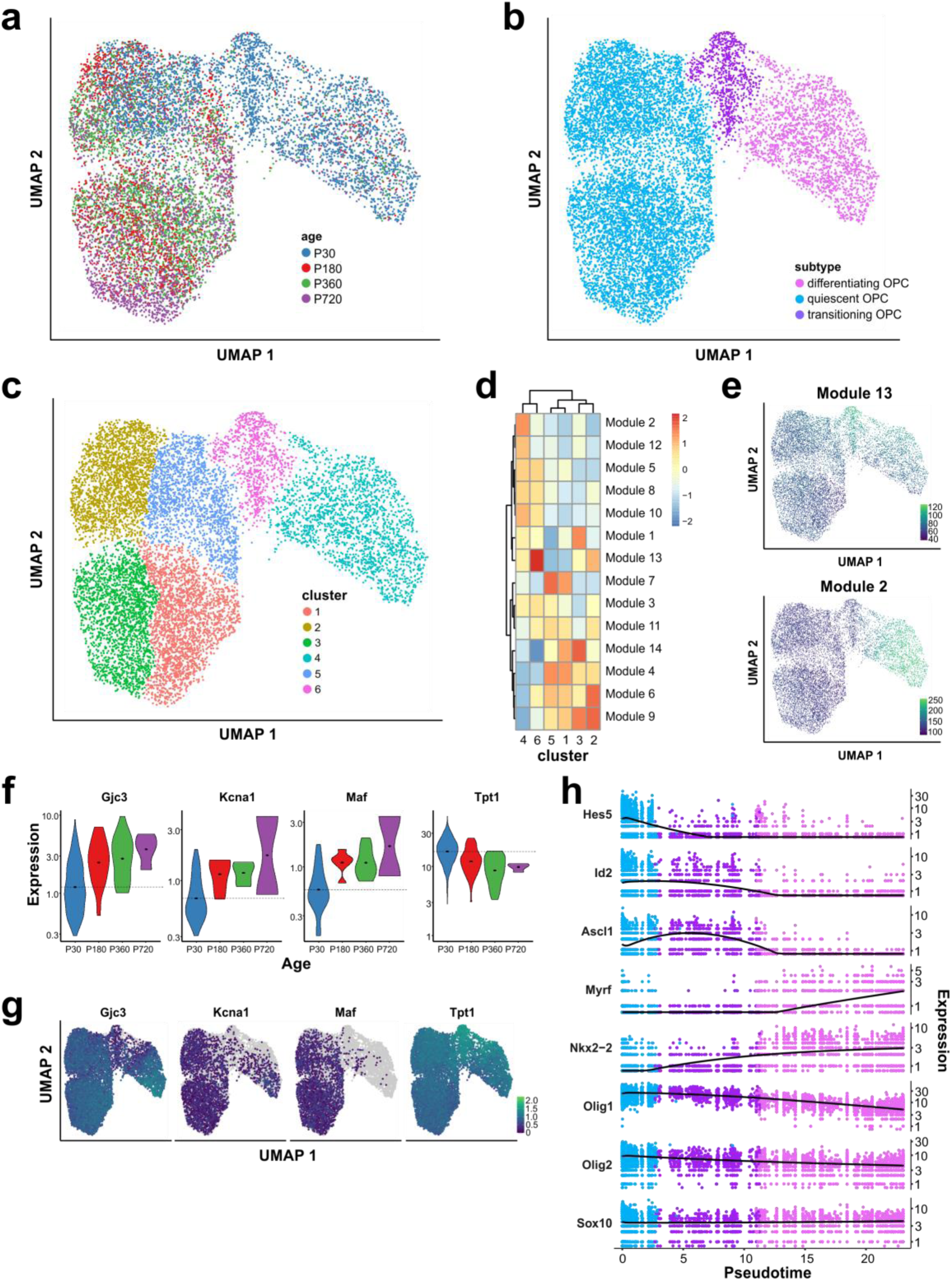
Identification of a novel transitioning OPC population poised to undergo differentiation. **a.** UMAP plot of differentiating OPCs and quiescent OPCs that immediately antecede the differentiating population, colorized by age groups. **b.** A colorized by different OPC subtypes (*Differentiating OPC*, *Quiescent OPC*, and *Transitioning OPC*). **c.** A colorized by different clusters (1-6) where cluster 6 represents the transitioning OPC subtype. **d.** Identification of different modules that represent groups of genes that are co-regulated in different clusters. **e.** UMAP plots of module 13 and module 2 that represent *Transitioning OPC* and *Differentiating OPC* groups, respectively. **f.** Expression violin plots of the top four genes (*Gjc3*, *Kcna1, Maf,* and *Tpt1*) differentially expressed across aging in *Transitioning OPC* (monocle3 linear regression, q-value < 0.001). **g.** UMAP plots of the top four genes in g across the four different age groups. The three genes that are downregulated in *Transitioning OPC* (*Gjc3, Kcna1,* and *Maf*) are upregulated with aging whereas *Tpt1*, which is upregulated in *Transitioning OPC* is downregulated with aging. **h.** Expression of different oligodendrocyte-related transcription factors across the pseudotime originating from *Quiescent OPC* subtype.

To explore the influence of aging on this transition, we identified 54 genes that were significantly differentially expressed as a function of age in the *Transitioning* population (Monocle likelihood ratio test, 0.05% FDR); among these were *Gjc3*, *Kcna1*, *Maf*, and *Tpt1*. Moreover, genes that increased expression with aging (*Gjc3*, *Kcna1*, and *Maf*) were strongly downregulated in *Transitioning* OPCs, raising the possibility that they must be downregulated to enable lineage progression. *Gjc3* encodes connexin 29 (Cx29), a gap junction protein expressed by oligodendrocytes and implicated in axonal communication and possible potassium uptake^49,50^. *Kcna1* encodes a voltage-gated potassium channel, K_v_1.1, that opens in response to membrane depolarization. Interestingly, both *Gjc3* and *Kcna1* have previously been shown to be a target of miR-27a, a microRNA that regulates oligodendrocyte development and survival^51^. *Maf* is a leucine zipper-containing transcription factor that in myelinating Schwann cells acts downstream of Neuregulin1 (NRG1) to regulate cholesterol biosynthesis^52^. In contrast, mRNA for *Tpt1*, which encodes the tumor protein, translationally-controlled 1 (also known as *Trt, p21, or p23*), an inhibitor of cyclin-dependent kinase (CDK), decreased with aging, but was enriched in the *Transitioning* OPC population (Fig. 4g,h). Although widely used as a senescence marker, p21 signaling has also been shown to be required for OPC differentiation following growth arrest^53^, raising the possibility that p21 may play a cell cycle-independent role in OPCs that are undergoing differentiation.

To investigate the temporal regulation of different transcriptional networks, a list of transcription factors involved in oligodendrogenesis was curated and plotted along the pseudotime trajectory across *Quiescent, Transitioning,* and *Differentiating* OPC subtypes (Fig. 4f). The transcriptional expression of *Sox10* remained constant throughout differentiation (Moran’s I value = 0.018), whereas *Olig1* and *Olig2* progressively decreased in expression (Moran’s I values = 0.632 and 0.216, respectively). *Nkx2-2*, a key transcription factor involved in OPC differentiation^54^, increased expression from *Quiescent* to *Transitioning* OPCs (Moran’s I value = 0.248). *Hes5*, *Id2,* and *Ascl1*, which have been shown to repress OPC differentiation^55^, were downregulated in *Transitioning* OPCs (Moran’s I values = 0.372, 0.168, and 0.241, respectively), which may relieve inhibition of pro-differentiating genes, such as *Myrf*^48^ (Moran’s I value = 0.429). In addition, we used the Monocle 3 graph_test algorithm, which utilizes Moran’s I statistics, to unbiasedly determine which transcription factors change in expression along the pseudotime trajectory of OPC differentiation (Supplementary Data 4). With the ability to isolate OPCs and define the transcriptional phenotype of OPCs in this key transition state, we defined dynamic changes in the expression of transcription factors that may influence the successful transition from quiescent to differentiating OPCs and their aging-associated decline in differentiation potential.

### Immune and cell death pathways are activated in OPCs with aging

Although quiescent OPCs appear phenotypically homogeneous within the cerebral cortex, in terms of their highly ordered distribution, morphology, and cellular dynamics, these characteristics may fail to reveal underlying transcriptional differences that may influence their ability to undergo state transitions and respond to injury and disease. To explore whether such hidden diversity exists, we applied unbiased dimensionality reduction to identify patterns of gene co-regulation in *Quiescent* OPCs, which revealed that this population of OPCs undergoes an aging-dependent shift in transcriptional profile (Fig. 5a,b). Using an NMF-based regulatory pattern identification, we identified an aging-associated pattern of co-regulated genes in aged *Quiescent* OPCs (Fig. 5c). Among the statistically significantly differentially expressed genes (Supplementary Data 5), *C4b* (complement component 4B) was significantly upregulated in aged OPCs compared to young OPCs (Monocle likelihood ratio test, 0.05% FDR; Fig. 5d). Fluorescence *in situ* hybridization in coronal brain sections from young and old *Matn4-mEGFP* mice confirmed that *C4b* mRNA is more abundant in GFP+ *Pdgfra+* OPCs in the aged brain (Extended Data Fig. 6a). As a key component of the complement cascade, *C4b* is involved in opsonizing target cells for removal by professional phagocytes. *C4b* has previously been shown to be significantly upregulated in OPCs^56^ and astrocytes^2^ with aging, as well as in mature oligodendrocytes in a mouse model of AD^57^.

**Figure 5.**
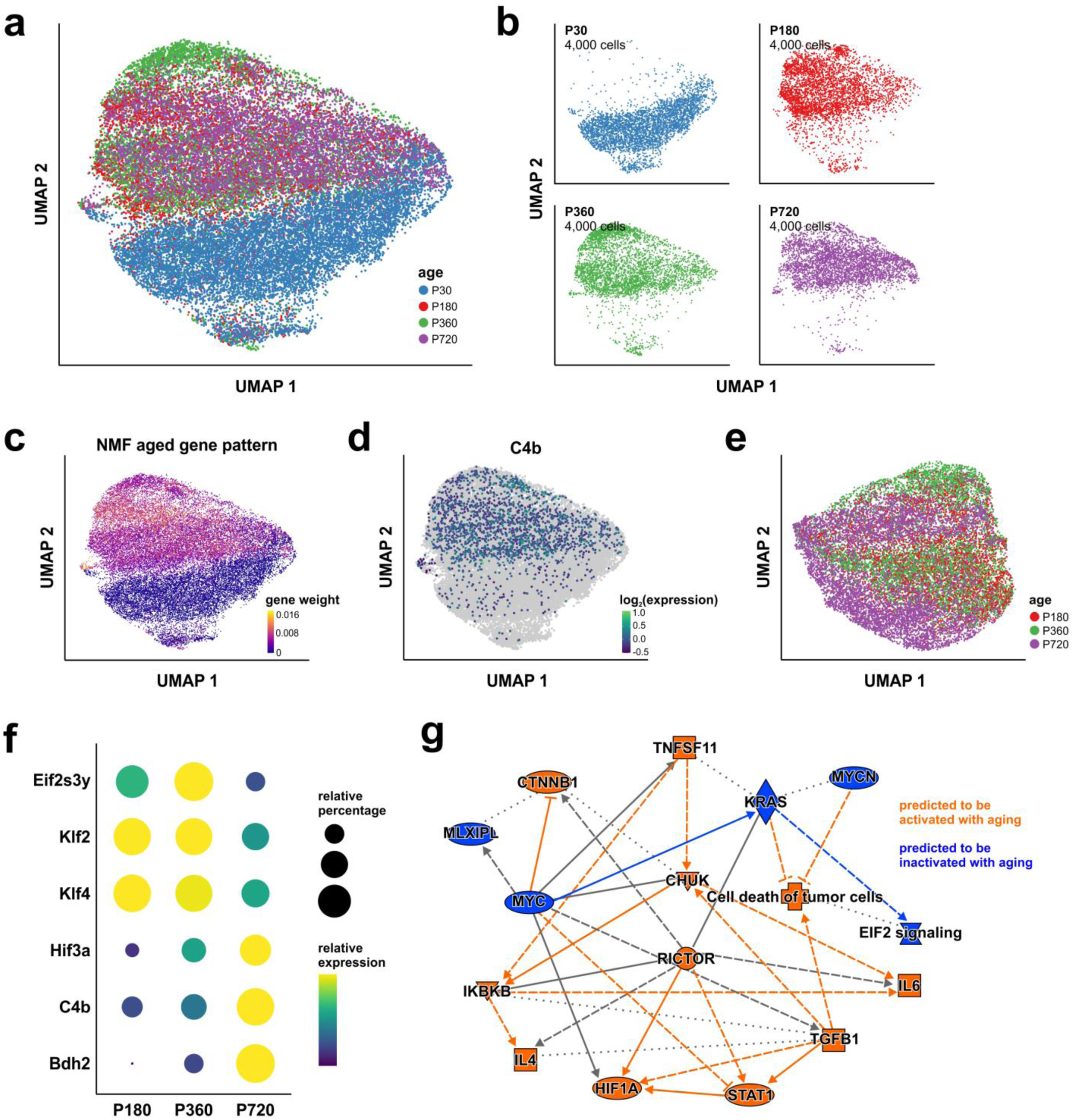
Quiescent OPCs undergo aging-associated transcriptional changes. **a.** UMAP plot of *Quiescent OPC* subtype, colorized by age groups. **b.** Separate UMAP plots for different age groups to show the separation of young, P30 quiescent OPCs from aged quiescent OPCs from P180, P360, and P720. 4,000 cells from each age group were randomly selected and plotted. **c.** Identification of an aged gene pattern that is associated with aged quiescent OPCs using Nonnegative Matrix Factorization (NMF). **d.** UMAP plot of *C4b* gene expression that shows the enrichment of expression in aged quiescent OPCs. **e.** UMAP of quiescent OPCs from P180-P720 brains. **f.** Examples of significantly differentially expressed genes with aging plotted in a dot plot where the relative percentage of cells is represented by the size of circles and relative expression is represented by color. **g.** Ingenuity Pathway Analysis (IPA) of statistically significantly differentially expressed genes in quiescent OPCs with aging shows that immune-pathways are predicted to be activated (orange) whereas cell growth pathways are predicted to be inactivated (blue).

To distinguish development from true aging, we removed young P30 cells from the dataset and reanalyzed the older P180-720 cells to test whether there still was a shift in the OPC transcriptome with aging (Fig. 5e). Of the top differentially expressed genes with respect to aging in *Quiescent OPC* from P180-720 animals, genes that encode Krüppel-like factors (KLF), *Klf2* and *Klf4*, and *Eif2×3y* were significantly downregulated in *Quiescent OPC* from the aged, P720 mouse cortex (Monocle likelihood ratio test, 0.05% FDR; Fig. 5f). Although KLF2 and KLF4 have been shown previously to regulate endothelial cell survival and function^58^, they have not been studied in oligodendrocyte lineage cells. In addition to *C4b*, *Hif*3a, which encodes hypoxia-inducible factor 3a, and *Bdh2*, which encodes 3-hydroxybutyrate dehydrogenase 2, were significantly upregulated in *Quiescent OPC* from P720 animals (Fig. 5f). *Hif3a* has been shown to be enriched in oligodendrocytes from experimental autoimmune encephalomyelitis (EAE) spinal cords^14^. BDH2 has been shown to be involved in reactive oxygen species (ROS)-induced cell death and autophagy in the context of cancer^59^. To investigate the upstream biological processes that may drive this striking transcriptional shift in *Quiescent OPC* with aging, we used Ingenuity Pathway Analysis (IPA) to identify potential upstream regulators^60^. The analysis revealed that immune and cell death pathways, including STAT1 and TGF-β1, were predicted to be activated in aged OPCs. A previous bulk RNA-seq analysis of young and aged rat OPCs has reported that EIF2 and IL-6 signaling pathways are enriched in aged OPCs^12^. We also found that pathways involved in cell growth, including MYC and KRAS, were predicted to be suppressed with age in OPCs (Fig. 5g). In line with our observation that MYC pathway is inhibited in aged OPCs, exogenous application of c-Myc has been shown to strongly rejuvenate aged OPCs and increase their proliferation and differentiation *in vitro*^61^. Together, our findings demonstrate that OPCs undergo significant aging-associated transcriptional changes, which may help identify potential targets to improve regeneration of oligodendrocytes in the aged CNS.

### Inhibition of HIF-1a and Wnt pathways promotes OPC differentiation *in vitro*

IPA upstream analysis of *Quiescent* OPCs indicated that HIF-1α and Wnt/β-Catenin signaling pathways are predicted to become more pronounced in these progenitors as the brain ages (Fig. 5g). Indeed, *Hif1a* expression in *Quiescent* OPCs increased progressively with aging (Fig. 6a, Extended Data Fig. 6b), which was also observed *in situ* at the protein level in the aged mouse cortex (Fig. 6b,c). To determine if inhibition of HIF- 1α pathway cell autonomously restores the differentiation potential of aged OPCs, we performed pharmacological manipulations in primary OPC cultures from young adult (YA) and aged adult (AA) mice^12^. OPCs acutely isolated from AA mice exhibited higher immunoreactivity to HIF-1α and reduced differentiation potential (Fig. 6d-e). When OPCs from AA mice were exposed to the HIF-1α inhibitor CAY10585 (Fig. 6f), the differentiation potential of aged OPCs was restored to that of young adult OPCs (Fig. 6g; Wilcoxon rank sum test with the Holm-Šídák multiple comparisons test). These results suggest that the increased expression and activation of HIF-1α by OPCs in the aged brain may directly impair OPC differentiation.

**Figure 6.**
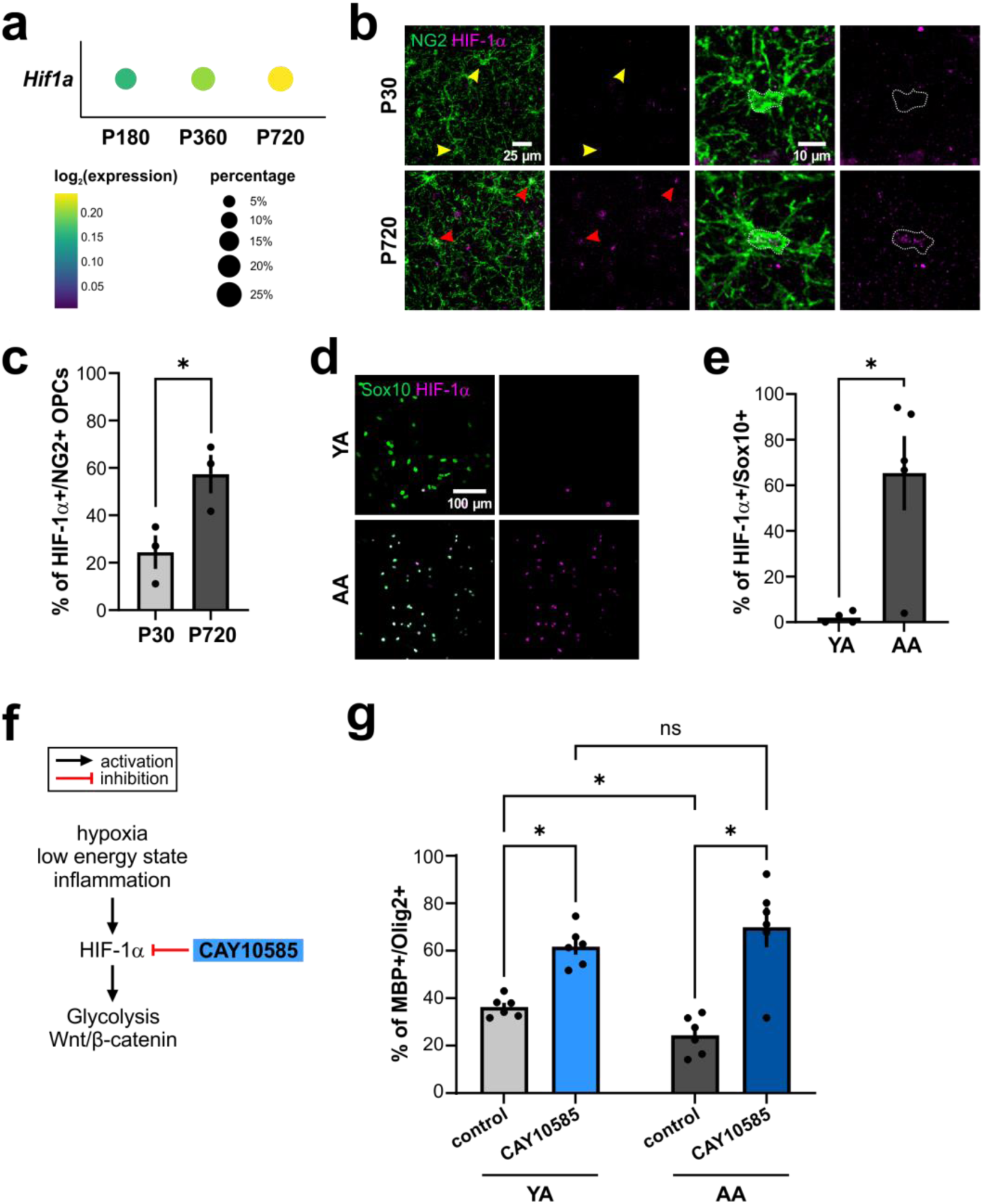
HIF-1α pathway is activated in aged OPCs, which functionally inhibits their differentiation. **a.** Dot plot of *Hif1a* expression in *Quiescent OPC* from P180, P360, and P720 timepoints. **b.** Immunofluorescence (IF) of HIF-1α (magenta) and NG2 (green) in P30 and P720 cortices. Yellow arrowheads indicate P30 NG2+ OPCs that lack HIF-1α immunoreactivity and red arrowheads indicate P720 NG2+ OPCs that show HIF-1α immunoreactivity. **c.** Quantification of the percentage of HIF-1α+ OPCs in young (P30) vs aged (P720) cortex (Student’s t-test, * p-value < 0.05). **d.** IF staining for HIF-1α (magenta) and Sox10 (green) in mouse OPC primary cultures. YA: young adult, AA: aged adult. **e.** Quantification of HIF-1α+ Sox10+ cells (Wilcoxon rank sum test, * p-value < 0.05). **f.** Schematic of CAY10585 drug impinging on the HIF-1α pathway. Hypoxia, low energy, and inflammation have been shown to activate HIF-1α signaling. At downstream, HIF-1α is involved in upregulating glycolysis and Wnt signaling. **g.** Quantification of MBP+ differentiating OPC/Olig2+ oligodendrocyte proportions with and without CAY10585 treatment in OPCs isolated from YA or AA (Wilcoxon rank sum test with the Holm-Šídák multiple comparisons test, * p-value < 0.05).

Previous studies have reported that hypoxia and the activation of HIF-1α can upregulate the downstream Wnt/β-Catenin signaling pathway in cancer ^62,63^. Given the increased expression of *Ctnnb1*, which encodes β-catenin, in *Quiescent* OPCs from aged mice (Extended Data Fig. 6c, Extended Data Fig. 7a), we tested whether the pharmacological inhibition of Wnt signaling in aged OPCs can recapitulate the effect of HIF- 1α inhibition. We used two different Wnt inhibitors to either globally block all Wnt signaling (IWP-2) or selectively attenuate the canonical Wnt pathway by promoting Axin stabilization (XAV939) (Extended Data Fig. 7b). Previously, XAV939 has been shown to increase myelination in both *ex vivo* mouse cerebellar slices and *in vivo* adult mouse spinal cords following lysolecithin-mediated demyelination^64^. We found that both IWP-2 and XAV939 exposure significantly increased the differentiation of OPCs (Extended Data Fig. 7c). Notably, the effects of these inhibitors were more pronounced in OPCs isolated from AA compared to those from YA mice (Extended Data Fig. 7c), suggesting that enhanced activation of canonical and non-canonical Wnt signaling pathways may contribute to the lower rate of OPC differentiation in the aged brain. Together, these studies highlight the ability of transcriptional profiling from *Matn4-mEGFP* mice to uncover changes in regulatory pathways critical for lineage progression in oligodendrocyte progenitors and that aging-induced mechanisms that reduce OPC differentiation may be pharmacologically reversible.

## Discussion

OPCs comprise a persistent, highly dynamic population of glial cells that remain widely distributed in the adult CNS^24^. In the aging brain, OPCs have been shown to upregulate senescence pathways^65^, engage in antigen presentation^13^, and associate with Aβ plaques^66^, suggesting that they may regulate inflammation and the extracellular environment; however, because they represent only a small fraction of all brain cells, aging dependent changes in their properties have been difficult to define, and the molecular mechanisms that govern their distinct behaviors and lineage progression remain poorly understood. To overcome these challenges, we developed *Matn4*-*mEGFP* mice, which exhibit restricted expression of membrane-anchored EGFP by OPCs throughout most areas of the CNS, which we then used to generate single-cell transcriptional profiles of large populations of acutely isolated OPCs from the cerebral cortex across the lifespan, defining the extent of their heterogeneity and delineating aging-associated deficits in key pathways that regulate their ability to differentiate into myelin-forming oligodendrocytes.

Genetic interrogation of OPCs is complicated by their dynamic nature, as they exhibit robust homeostasis to keep their density in the neuropil constant, dividing when members of their population die, differentiate, or transform into scar/barrier cells after injury^24,67,68^. OPCs also retain the ability to mature into oligodendrocytes when conditions are appropriate for new myelin formation, a process that continues in the adult CNS and can be enhanced by increased neuronal activity and environmental enrichment^9,69^. Thus, extensive OPC sampling is required to resolve the transcriptional changes associated with these state transitions and determine what proportion of these cells are mobilized in different contexts. Previous efforts to define the diversity of OPCs and their underlying transcriptional heterogeneity have been limited by the low capture rate of these cells using bulk isolation procedures^25,70,71^. The >38,000 OPCs represented in our scRNA-seq dataset provide additional ground-truth OPC transcriptomic data to populate these transitional states, define their heterogeneity, and assist in the identification of not only aging-dependent transcriptional changes but also those associated with key state transitions such as proliferation and differentiation. We show that these data can be projected onto human OPCs obtained through bulk isolation, highlighting conservation of the mechanisms responsible for OPC behavior and the value of this high-resolution dataset for revealing reveal transitional states in other contexts.

*In vivo* fate tracing studies suggest that oligodendrocytes in both the brain and spinal cord arise in temporarily distinct waves from different ventral and dorsal regions of the ventricular germinal zones during development^72–74^. In the cerebral cortex, ventrally derived OPCs are eventually replaced by a dorsally derived pool^72^. Consistent with this replacement hypothesis and a previous developmental OPC transcriptomics study^71^, our studies show that *Quiescent* OPCs in the cortical gray matter have a remarkably consistent transcriptional profile, and there was little evidence of distinct populations of OPCs specialized for different functions. Indeed, the largest deviations from the *Quiescent* pool were associated with either cell state change or aging (Fig. 2b,c; Fig. 5a,b). Although we did not identify clear diversification within cortical OPCs, it is possible that oligodendrocyte lineage cells exhibit strong regional differences within the brain^75–77^. Recently, large-scale transcriptomic profiling of the marmoset CNS showed that OPCs, along with other glial cells, exhibit strong regional diversity, with white matter and gray matter OPCs clustering in distinct groups according to transcriptional features^78^. In support of this conclusion, OPCs in white matter have been shown to exhibit a higher proliferation rate than those in gray matter and exhibit distinct electrophysiological properties^5,32,33,79^. Moreover, in the developing zebrafish spinal cord, two populations of OPCs have been observed, which vary in location, activity patterns, and differentiation rates^80^.

Our data support the conclusion that even in the aged brain (P360-720), OPCs retain their ability to enter the cell cycle and self-renew (Fig. 2e), consistent with *in vivo* imaging studies of OPCs demonstrating occasional cell division^24^, evidence of thymidine analog incorporation^31,81^, expansion of genetically traced OPC clones^32^, and the maintenance of OPC density across ages despite ongoing oligodendrogenesis^82^. However, by six months of age, the proportion of cells engaged in these dynamic behaviors was small (∼17%; Fig. 2e), consistent with the decrease in oligodendrogenesis observed through fate tracing studies^31,83^. Thus, the decline in proliferation may simply reflect that decline in the production of new oligodendrocytes, rather than a cell-intrinsic change in the ability of the cells to proliferate. In support of this conclusion, focal laser ablation^24^ or genetic ablation of OPCs^84^ enhances the proliferation of nearby OPCs in the adult CNS.

In addition to their role as oligodendrocyte progenitors, some OPCs engage in structural remodeling of neurons through engulfment^85–87^, and they migrate to sites of focal injury much like microglia, contributing to the formation of glial barriers/scars^68,88^. Expression of chondroitin sulfate proteoglycans by OPCs has been shown to limit axon regrowth in the spinal cord^89^, indicating that this transformation has a critical impact on recovery processes. However, it has been difficult to track the transformation of quiescent OPCs into this reactive state, as classic genetic markers of OPCs (*Cspg4*, *Pdgfra*, and *Olig2*) no longer become restricted to OPCs following injury^90^. Our results indicate that *Matn4-*mEGFP transgene expression remains faithful to OPCs even following traumatic injury to the brain (Extended Data Fig. 8), suggesting that these mice will be useful for studying the molecular basis for distinct OPC behaviors and reveal their contributions to tissue repair.

In the EAE model of MS in rodents, as well as in human MS patient tissue samples, a subset of oligodendrocyte lineage cells has been shown to exhibit immunological profiles, characterized by the upregulation of MHC class I and II^14,91,92^. It is thought that these immune-associated transcriptional changes are induced by the release of interferon-γ (IFN-γ) and other cytokines by CD8+ T cells and microglia^93,94^. Consistent with recent studies of oligodendroglia in MHC class I and II reporter mice^95^, we did not detect significant upregulation of these pathways in these naïve mice, suggesting that aging-related increases in overall inflammatory state alone are not sufficient to induce this phenotypic change^65^. Previous studies have also proposed that some OPCs exhibit senescent features in aging and disease conditions, in which OPCs have decreased potential for differentiation and oligodendrocyte regeneration due to increased expression of p16 and p21^66,96^. Indeed, our scRNA-seq data predict that aging is associated with higher activation of immune and cell death pathways, including STAT1 and TGF-β1, whereas pathways involved in cell growth, including MYC and KRAS, are predicted to be suppressed (Fig. 5g). Myc overexpression has previously been shown to promote functional rejuvenation of aged OPCs, while its inhibition in neonatal OPCs induced an aged-like phenotype^61^. Here, we identified additional pathways, HIF-1α and Wnt, that are enhanced in aged OPCs and inhibit their differentiation capacity. Previous *ex vivo* and *in vitro* studies have shown that HIF-1α signaling may activate the canonical Wnt pathway or regulate Sox10 to inhibit OPC differentiation and myelination^97,98^. Together, these results suggest that an oligodendrocyte lineage-specific inhibition of HIF-1α and Wnt pathways may provide a potential therapeutic avenue to promote the regeneration of oligodendrocytes and remyelination in aging and disease, where local OPCs exhibit reduced differentiation^99^.

## Online Methods

### Animal care and use

Female and male adult *Matn4-mEGFP* mice were used for experiments and randomly assigned to experimental age groups. Mice were maintained on a 12-hour light/dark cycle, housed in groups no larger than 5, and food and water were provided *ad libitum*. All animal experiments were conducted in accordance with the National Institute of Health Guide for the Care and Use of Laboratory Animals and were approved by Animal Care and Use Committee at Johns Hopkins University (Protocol numbers: MO23M202, MO20M344).

### Generation of Matn4-mEGFP mouse line

*Matn4-mEGFP* (*Matn4^mEGFP/+^*) mouse line was generated by knocking in MARCKS (ATGGGTTGCTGTTTCTCCAAGACC), EGFP, WPRE, and bGH-polyA sequences into the first coding exon of *Matn4* using CRISPR-Cas9 with 800 bp homology arms. Whole-genome sequencing was performed to ensure a single insertion into the correct genetic locus. *Matn4-mEGFP* mice were genotyped using polymerase chain reaction (PCR) analysis of DNA isolated from toe snips taken at postnatal day 5 (P5). The wild-type allele was identified by a 178 bp PCR product and the mutant, knock-in allele by a 356 bp PCR product using the following primers: MATN4-mEGFP-WT-F (ACACTGTGGTTCGTCATCCT), MATN4-mEGFP-WT-R (accctggctcactgtggata), MATN4-mEGFP-KI-R (AAGAAGATGGTGCGCTCCT). Reactions were run under the following conditions: 95°C × 3 min, (95°C × 30 s, 63°C × 30 s, 72°C × 60 s) × 35 cycles, 72°C × 7 min using the KAPA Express PCR kit (Extended Data Fig. 1b).

### Immunohistochemistry

Mice were deeply anesthetized with the i.p. injection of pentobarbital (100 mg/kg) and perfused transcardially with 20 mL of 0.1 M phosphate buffered saline (1x PBS) and then 20 mL of freshly prepared, ice-cold 4% paraformaldehyde (PFA, Electron Microscopy Sciences, #19210) in 1x PBS (pH7.4). Dissected tissues were post-fixed in 4% PFA/PBS at 4°C in dark for 4 hours, and then cryoprotected in 30% sucrose/0.1 % sodium azide in 1x PBS at 4°C in dark for at least 48 hrs. Before collecting free-floating sections, tissue samples were embedded in Tissue-Tek O.C.T Compound (Sakura Finetek, #4583) and sectioned at −20°C using a Thermo Scientific Microm HM 550 at the thickness of 35 μm. Before immunostaining, sections were rinsed briefly in PBS and then permeabilized with 0.5% Triton X-100 in 1x PBS for 10 min at room temperature (RT). To prevent non-specific binding of antibodies, brain sections were further incubated in the blocking buffer (10% normal donkey serum, Jackson Immuno, #017-000-121, and 0.3% Triton X-100 in 1x PBS) for 1 hour at RT, followed by the primary antibody incubation at RT overnight. After washed in 1x PBS for 3 times, 10 min each, brain sections were then incubated with the secondary antibodies for 2 hours at RT before another wash in 1x PBS as described above. Both primary and secondary antibodies were diluted in the blocking buffer. Sections were mounted on slides with Aqua-Poly/Mount (Polysciences, #18606). Images were acquired using Zeiss LSM 800 and 880 confocal microscopes and analyzed using ImageJ (https://imagej.net/software/fiji/). Primary antibodies used in this study: guinea pig anti-NG2 (Bergles lab, 1: 5000), rabbit anti-PDGFRa (Cell Signaling, 1:1000), chicken anti-GFP (Aves, #GFP-1020, 1:4000), rabbit anti-HIF1a (Novus Biologicals, NB100-479, 1:200; for IHC), rabbit anti-Olig2 (Millipore, AB9610), goat anti-Sox10 (R&D Systems, AF2864), rat anti-MBP (Bio-rad, MCA409S), and rabbit anti-HIF1a (Abcam, AB179483; for *in vitro* IF). Secondary antibodies used for mouse brain IHC were all purchased from Jackson ImmunoResearch and used at 1:1000: Cy3 donkey anti-guinea pig IgG (#706-165-148), Alexa Fluor 647 donkey anti-rabbit IgG (#711-605-152), and Alexa Fluor 488 donkey anti-chicken IgG (#703-546-155). Alexa Fluor dye-conjugated secondary antibodies used for *in vitro* IF were purchased from Invitrogen.

### Head plate installation and cranial window surgery

Cranial windows were prepared as previously described^100^. Briefly, mice were anesthetized with inhaled isoflurane (0.25-5%) and placed in a customized stereotaxic frame. Surgery was performed under standard and sterile conditions. After hair removal and lidocaine application (1%, VetOne, NDC 13985-222-04), the mouse’s skull surrounding the right motor cortex was exposed and the connective tissue was carefully removed from the skull. Vetbond™ (3M) was used to close the incision site. A custom-made metal head plate was fixed to the cleaned skull using dental cement (C&B Metabond, Parkell Inc.). A piece of the cover glass (VWR, No. 1) was placed in the craniotomy and sealed with cyanoacrylate glue (VetBond (3 M) and Krazy Glue). Animals were allowed to recover in their home cages for at least 2 weeks before being subjected to imaging.

### In vivo two-photon laser scanning microscopy

Two-photon laser scanning microscopy was performed with a Zeiss LSM 710 microscope equipped with a GaAsP detector using a mode-locked Ti-Sapphire laser (Coherent Chameleon Ultra II) tuned to 920 nm. The head of the mouse was immobilized by attaching the head plate to a custom machined stage mounted on the microscope table. Fluorescence images were collected 50-200 μm from the cortical surface using a coverslip-corrected Zeiss 20x/1.0 W Plan-Apochromat objective.

### Cell Isolation, Enrichment, and cDNA Library Preparation

Single-cell suspension of *Matn4-mEGFP* mouse cortical OPCs was achieved using the Miltenyi Neural Tissue Dissociation Kit (Miltenyi Biotec 130-092-628) followed by fluorescence-activated cell sorting (FACS) isolation of GFP+ cells with BD FACS Aria IIu Cell Sorter (BD Biosciences) through the Flow Cytometry Core at Johns Hopkins Medicine. Briefly, the mice were anesthetized with the i.p. injection of pentobarbital (100 mg/kg) and perfused transcardially with 20 mL of Hanks’ Buffered Salt Solution lacking Mg^2+^ and Ca^2+^ (HBSS–) (Gibco). The dissected cortices were coarsely chopped in HBSS– on ice and centrifuged at 300 x g for 2 min at RT. Subsequent enzymatic dissociation of the tissue was performed based on the manufacturer’s protocol (Miltenyi Biotec 130-092-628). Debris Removal Solution (Miltenyi Biotec 130-109-398) was used to effectively remove cell and myelin debris. Cells were then resuspended in 1% FBS/HBSS–/2 mM EDTA/25 mM HEPES buffer for FACS. The isolated cells were collected in EDTA-free buffer (1% FBS/HBSS–/25 mM HEPES) and spun down in LoBind 1.5 mL tubes (Eppendorf 022431081) to make ∼1,000 cells/μl and processed using 10x Genomics Chromium Single Cell 3’ v3 and v3.1 kits according to the manufacturer’s protocol (10x Genomics). The libraries were QC-ed using the BioAnalyzer before being pooled and sequenced on the Illumina NovaSeq 6000 at ∼50,000 reads/cell (estimated from the initial loading).

### Data analysis

Following sequencing, data were pseudoaligned to the publicly available mouse reference genome using the kallisto-bustools pipeline^101^ to generate a cellxgene matrix for each biological sample. Downstream analyses were performed following the Monocle 3 pipeline^102^. The expression dataset was log-normalized (with a pseudo-count of 1) and the lower dimensional space was calculated using principal component analysis (PCA). Batch effects were corrected using the mutual nearest neighbor algorithm as described for visualization purposes^103^. The Uniform Manifold Approximation and Projection (UMAP) algorithm was used for two-dimensional reduction of the data^104^. Community detection was performed using the Monocle 3 cluster_cells method, based on Louvain/Leiden community detection with default settings. To identify transcript expression modules within the clusters or subtypes of interest, we used the Monocle 3 graph_test algorithm (monocle3::graph_test) that implements Moran’s I statistics to identify pattern of expression in a two-dimensional reduced expression data. To test for differences in gene expression, the Monocle 3 implementation of regression analysis (monocle3::fit_models) was used. For human OPC snRNA-seq dataset analysis, the pre-curated OPC dataset was downloaded from CZI cellxgene discover collections (supercluster: oligodendrocyte precursor): https://cellxgene.cziscience.com/collections/283d65eb-dd53-496d-adb7-7570c7caa443. OPCs from cerebral cortex were isolated and reprocessed using the Monocle 3 pipeline as detailed above. Mouse gene patterns associated with different OPC subtypes were defined using scCoGAPS^35^, and ProjectR^36^ was used to project those mouse patterns onto the human dataset following instructions from projectR Vignette.

### Immunofluorescence and RNA fluorescence in situ hybridization chain reaction (HCR IF + HCR RNA-FISH)

Tissue was perfused and dehydrated as described above for immunofluorescence, then sectioned at 16 μm onto slides. These slides were immediately placed at −20°C for one overnight, then stored at −80°C. HCR IF + HCR RNA was performed using the manufacturer’s protocol (Molecular Instruments; Schwarzkopf *et al*., 2021). EGFP signal in the *Matn4^mEGFP/+^*tissue was detected using chicken anti-GFP (Aves, #GFP-1020, 1:4000) followed by a Donkey Anti-Chicken B5 secondary antibody probe (Molecular Instruments). Probes targeting *Pdgfra*, *C4b, Hif1a,* and *Ctnnb1* were designed and purchased through Molecular Instruments. These probes were all amplified using hairpin amplifies also purchased through Molecular Instruments. Following the HCR IF + HCR RNA-FISH protocol, lipofuscin was quenched using TrueBlack Plus (1:40 in 1xPBS, Biotium) for 2 min, washed twice by immersing slides in 1x PBS for 5 min, and stained with DAPI (1:5,000 in 1x PBS, BioLegend) for 10 minutes. Slices were mounted using Aqua-Poly/Mount (Polysciences, #18606), and images were acquired using Zeiss LSM 880 confocal microscopes. All images were taken in layers 1 to 3 of the motor cortex.

### Image analysis of FISH

HCR IF + HCR RNA-FISH signal quantification was performed using Imaris x64 v9.9.1 (Bitplane). Cells robustly positive for both GFP and *Pdgfra* signal were included in the analysis. Satellite cells, actively dividing cells, and pyknotic cells were excluded from analysis, and ROIs were drawn around healthy, individual OPCs. Using the Surfaces tool, a volume representing the OPC soma was generated based on *Pdgfra* signal intensity. The Mask tool was then used to select only signal within this volume, and HCR puncta for each gene of interest was counted with the Spots tool, with a uniform intensity threshold applied across all samples for each gene of interest.

### Isolation of oligodendrocyte progenitor cells

Mouse OPCs were isolated similarly to rat OPCs with smaller modifications (treating two mouse brains as one rat brain) using a protocol previously described in detail^12^. In short, telencephalon and cerebellum were dissected in ice-cold isolation medium (Hibernate-A, Brainbits). The tissue was minced into 1-mm^3^ pieces and washed in HBSS– (Gibco). Adult mouse brain was mixed with 5 ml of dissociation solution (34 U/ml papain (Worthington) and 20 μg/ml DNAse Type IV (Gibco) in isolation medium). Brain tissue was dissociated on a shaker (50 r.p.m.) for 40 min at 35 °C. Digestion was stopped by the addition of ice-cold HBSS–. To obtain a single-cell suspension, the tissue was triturated in isolation medium supplemented with 2% B27 and 2 mM sodium pyruvate, first using a 5-ml serological pipette and then three fire-polished glass pipettes (opening diameter >0.5 mm). The supernatant containing the cells was filtered through 70-μm strainers into a tube containing 90% isotonic Percoll (GE Healthcare, no. 17-0891-01, in 10 × PBS pH 7.2 (Lifetech) after each round of trituration. The solution was topped up with DMEM/F12 (Gibco) and mixed to yield a homogenous suspension with a final Percoll concentration of 22.5%. The single-cell suspension was separated from remaining debris particles by centrifugation (800g, 20 min, room temperature, without a break). Myelin debris and all layers without cells were discarded, and the brain-cell-containing phase (final 2 ml) and cell pellet were washed in HBSS–. Red blood cell lysis buffer (Sigma, no. R7757) was used to remove blood cells. After centrifugation, cells were resuspended in 135 µl of MWBI and 15 µl of mouse FcR block solution (Miltenyi Biotec130-092-575). OPCs were isolated by positive selection for A2B5 using the MACS protocol according to the manufacturer’s instructions, MS columns (Miltenyi, no. 130-042-201) and MiniMACS Separators (Miltenyi, no. 130-042-102). 1.7 μl of mouse A2B5 IgM antibody (Millipore, MAB312) was used for two adult mouse brains; 20 μl of rat anti-mouse IgM antibody (Miltenyi, no. 130-047-302) was used per brain for magnetic labeling. A2B5-positive cells were flushed from the column with 1 ml of prewarmed CO2- and O2-pre-equilibrated OPC medium.

### Culture of adult oligodendrocyte progenitor cells

Isolated OPCs were seeded into 96-well plates (InVitro-Sciences) pre-coated with 5 μg/ml Poly-D-Lysine (Sigma) for 45 min at 37°C followed by wash off with dH_2_O containing 10 µg/ml Laminin (Fisher) for 2 hr. After isolation, OPCs were left to recover in 150 µl OPC medium (60 μg ml N-acetyl cysteine (Sigma), 10 μg/ml human recombinant insulin (Gibco), 1 mM sodium pyruvate (Gibco), 50 μg/ml apo-transferrin (Sigma), 16.1 μg/ml putrescine (Sigma), 40 ng/ml sodium selenite (Sigma), 60 ng/ml progesterone (Sigma), 330 μg/ml bovine serum albumin (Sigma) with 2% B27 (Gibco)), supplemented with basic fibroblast growth factor (bFGF) and platelet-derived growth factor (PDGF) (30 ng/ml each, Peprotech). OPCs were incubated at 37°C, 5% CO_2_ and 5% O_2_. The medium was completely exchanged for OPC medium with 20 ng/ml PDGF-AA and bFGF to remove dead cells. After 3 days, 50% of the cell culture medium was exchanged with fresh growth factors (OPC medium + 20 ng/ml bFGF and PDGF). On day 4, the entire medium was switched to promote further differentiation (OPCM + 40 ng/ml T3). The differentiation medium was replaced completely every 2-3 days with fresh growth factors or other small molecules were added fresh to the culture. Small molecules used in this study: 20 nM of XAV939 (Tocris, 3748), 1.2 μM of CAY10585 (Cayman Chemical, 10012682), and 0.4 μM IWP-2 (Tocris, 3533/10). Cells were fixed at day 7 of differentiation.

### Imaging and quantification of in vitro pharmacological assays

All images were acquired as single-plane images using an Opera High Content Screening System (Revvity). For imaging all areas adjacent to the edge of the well were omitted. To quantify the differentiation frequency of OPCs into oligodendrocytes we used Harmony High Content Imaging and Analysis Software (Revvity) to unbiasedly identify Olig2+ nuclei. From these nuclei, the software automatically identified MBP signal to segment potential oligodendrocyte cell bodies. We manually determined cut-off values for MBP+ oligodendrocytes based on median intensity measurement of the MBP signals in manually inspected random images. Based on these cut-off values, we labeled cells as differentiated oligodendrocytes. For statistical analyses, we performed Wilcoxon rank sum test with the Holm-Šídák multiple comparisons test.

## Data availability

All raw and preprocessed sequencing data generated for this study as well as the processed Monocle 3 cell_data_set (cds) object have been deposited in NCBI Gene Expression Omnibus (GEO) with accession code GSE249268. To promote open access to data, we also generated an interactive website to search the annotated dataset (https://tinyurl.com/aging-opcs).

## Acknowledgments

This research was supported by grants from the NIH (AG072305, NS041435), the Goldman Foundation, and the Dr. Miriam and Sheldon G. Adelson Medical Research Foundation. D.H. and A.K. were supported by fellowships from the NIH (F31NS110204 and F30AG084193, respectively). Y.M. was supported by a fellowship from the National MS Society (FG-1708-28962). We thank Chip Hawkins at JHMI Transgenic Core Laboratory for performing CRISPR/Cas9 microinjections and assisting in the generation of *Matn4-mEGFP* mouse line. We also thank Dr. Michele Pucak and Dr. Aleksandr Smirnov at JHMI Neuroscience Imaging Center for their assistance with image acquisition and analysis. We also thank our colleagues for their invaluable support throughout this study.

---

**Extended Data Figure 1.**
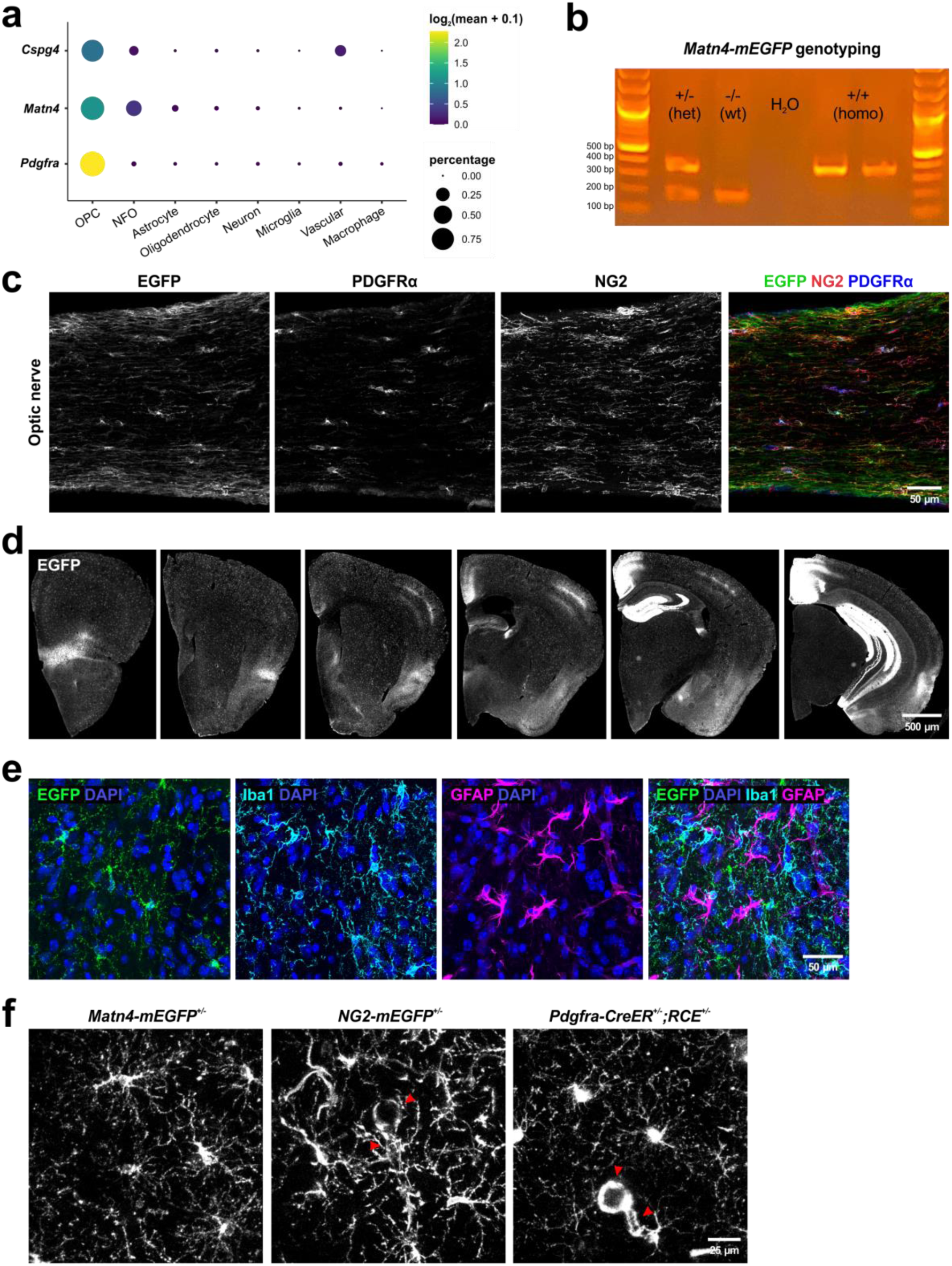
*Matn4-mEGFP* expression is restricted to OPCs and a subset of neurons. **a.** *Matn4* expression is specific to OPCs and newly-formed oligodendrocytes (NFO) in 6-7 week-old mouse V1 cortex (reanalysis of a publicly available scRNA-seq dataset^18^). **b.** Genotyping result of *Matn4-mEGFP* mouse line. Wildtype (wt) band size is 178 bp whereas the mutant, knock-in band size is 356 bp. **c.** EGFP signal in the optic nerve of *Matn4-mEGFP* mouse line is restricted to NG2+ PDGFRα+ OPCs. **d.** *Matn4-mEGFP* is also expressed by hippocampal granule cells and neurons in the somatosensory cortex barrel field and retrosplenial cortex. **e.** *Matn4-mEGFP* signal is absent from Iba1+ microglia and GFAP+ astrocytes. **f.** *In vivo* imaging of GFP+ cells in *Matn4-mEGFP*, *NG2-mEGFP*, and *Pdgfra-CreER; RCE* mouse lines. None of the vascular cells (red arrowheads) express EGFP in the cortex of *Matn4-mEGFP* mice.

**Extended Data Figure 2.**
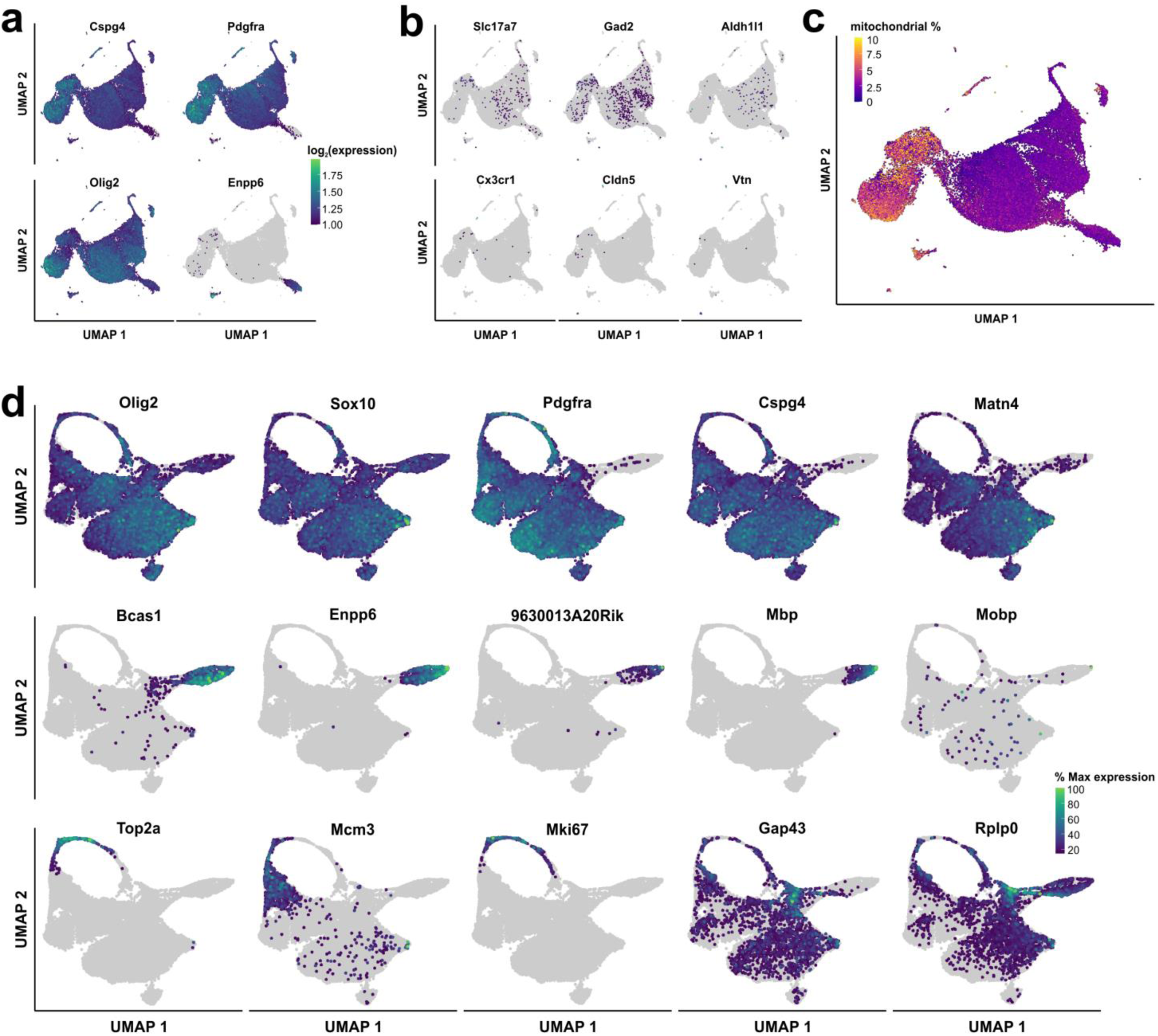
Preprocessing of the OPC scRNA-seq dataset. **a.** Expression of oligodendrocyte lineage cell genes in the uncleaned dataset. Most cells in the dataset express *Cspg4*, *Pdgfra*, and *Olig2* (OPCs) or *Enpp6* and *Olig2* (differentiating OPCs). **b.** Only a small group of cells that were FACS isolated from *Matn4-mEGFP* mouse line express non-oligodendrocyte lineage cell genes. **c.** UMAP plot of uncleaned dataset colorized by the percentage of mitochondrial-related genes (cutoff at 10%). Those cells with relatively high mitochondrial gene ratio (> 5%) were removed for downstream analyses. **d.** Expression of classic oligodendrocyte lineage marker genes (oligodendrocyte lineage: *Olig2, Sox10*; OPC: *Pdgfra, Cspg4, Matn4*; differentiating OPC: *Bcas1, Enpp6, 9630013A20Rik*; oligodendrocyte: *Mbp, Mobp*) as well as the subtype marker genes identified in this study (*Cycling OPC*: *Top2a, Mcm3, Mki67*; *Transitioning OPC*: *Gap43, Rplp0*) in the cleaned, preprocessed, final dataset.

**Extended Data Figure 3.**
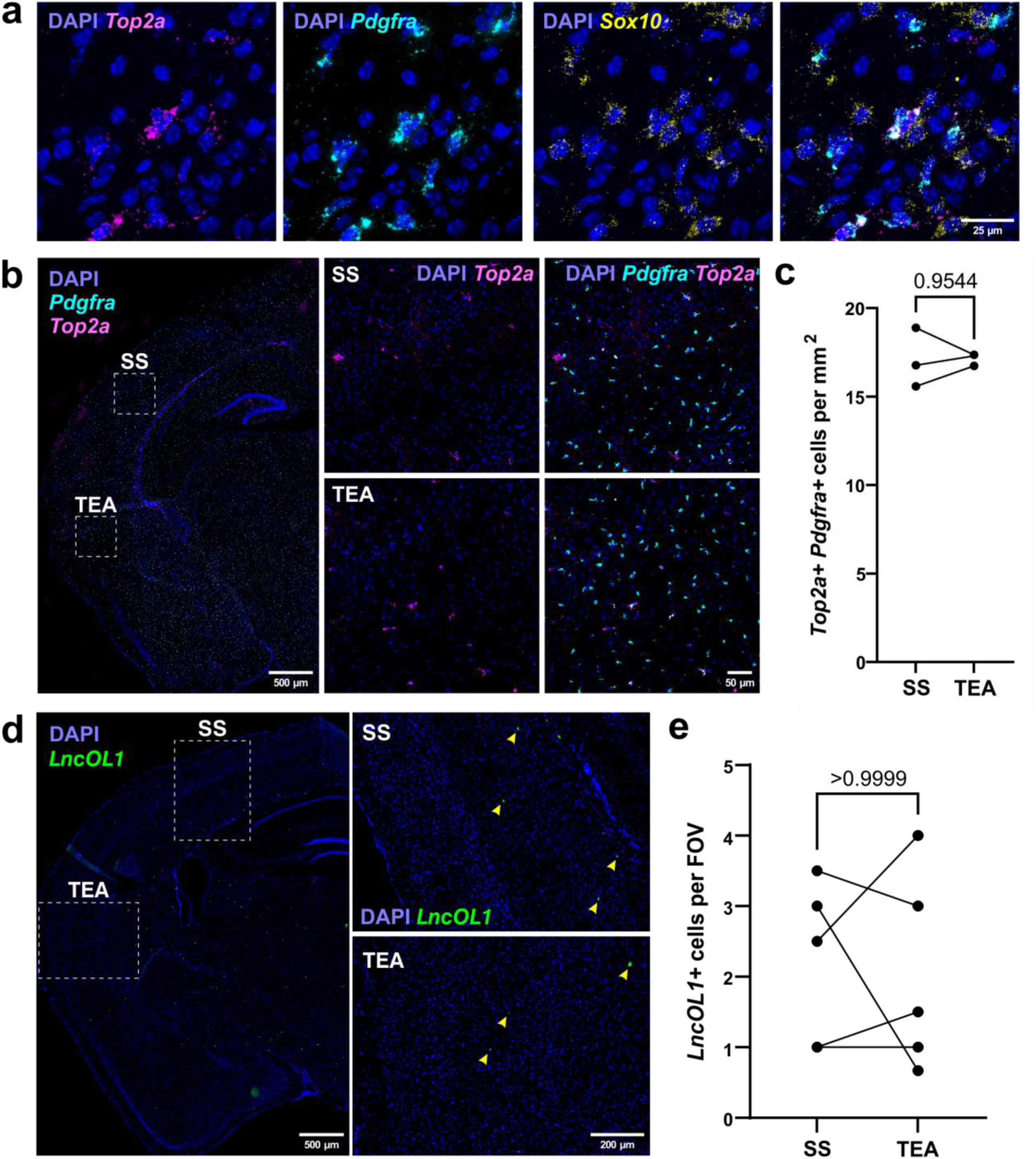
Fluorescent *in situ* hybridization (FISH) for *Cycling* and *Differentiating* OPC subtypes. **a.** FISH for *Top2a*, *Pdgfra*, and *Sox10* to identify *Cycling OPC in situ* in postnatal day 9 (P9) mouse brain. **b.** Comparison of the density of *Cycling OPC* in highly myelinated, somatosensory cortex (SS) and that in sparsely myelinated, temporal association cortex (TEA). **c.** Quantification of the density of *Cycling OPC* (*Top2a+ Pdgfra+*) in SS and TEA. **d.** FISH for *LncOL1* to identify *Differentiating OPC in situ* in P74 mouse brain. **e.** Quantification of the frequency of *LncOL1+ Differentiating OPC* in SS and TEA.

**Extended Data Figure 4.**
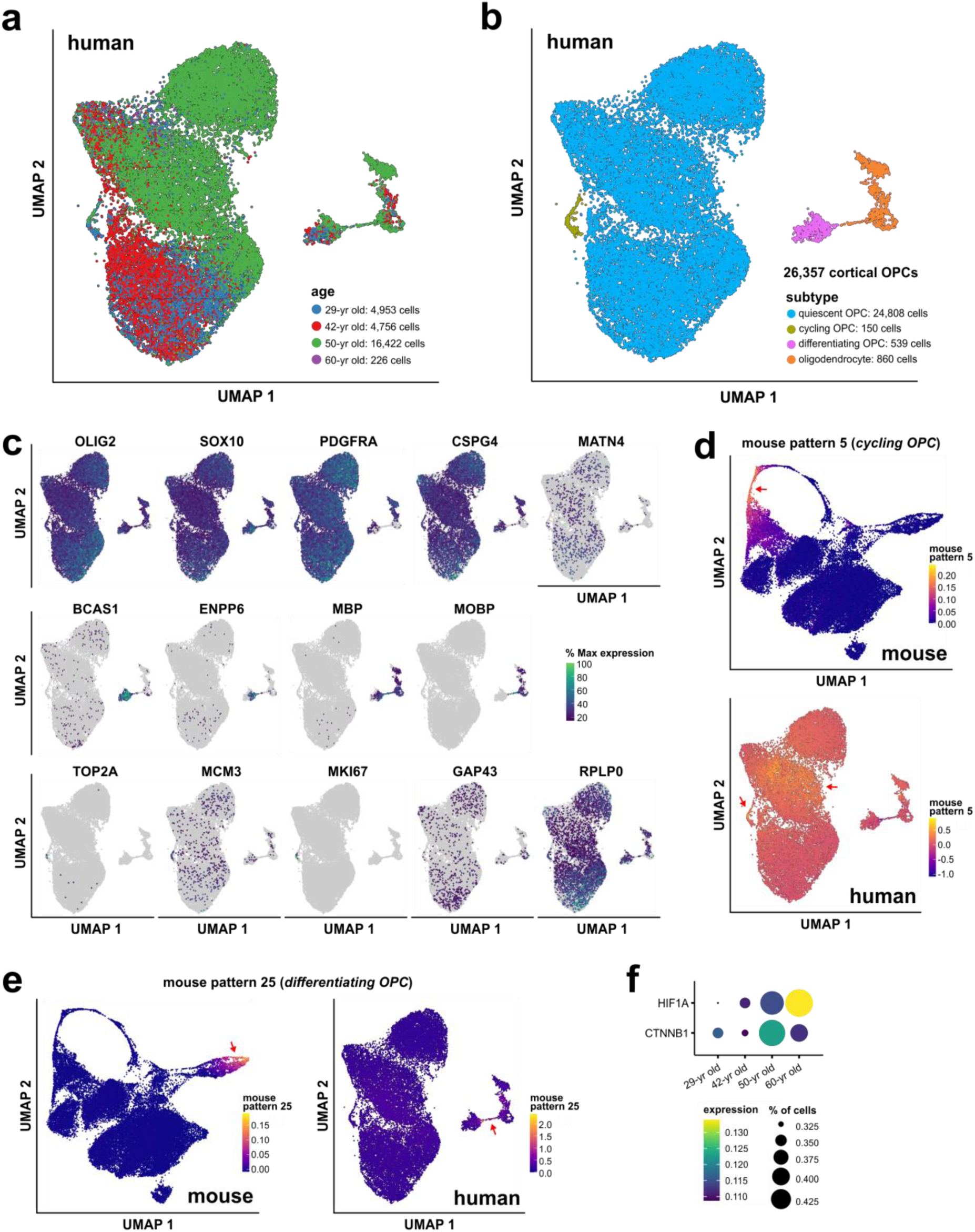
Reanalysis of human OPCs in Siletti *et al.* (2023) dataset^34^. **a.** UMAP plot of the dataset colorized by their four age groups (blue: 29-yr old, red: 42-yr old, green: 50-yr old, and purple: 60-yr old). **b.** UMAP plot of 26,357 human cortical OPCs colorized by their identified subtypes (quiescent OPC, cycling OPC, differentiating OPC, and oligodendrocytes). **c.** Expression of classic oligodendrocyte lineage marker genes (oligodendrocyte lineage: *OLIG2, SOX10*; OPC: *PDGFRA, CSPG4, MATN4*; differentiating OPC: *BCAS1, ENPP6*; oligodendrocyte: *MBP, MOBP*) as well as the subtype marker genes identified in this study *(Cycling OPC*: *TOP2A, MCM3, MKI67*; *Transitioning OPC*: *GAP43, RPLP0*) in the human cortical OPC dataset. **d.** UMAP plots of mouse *Cycling OPC* gene pattern 5 projected on the mouse scRNA-seq dataset and on the human snRNA-seq dataset. **e.** UMAP plots of mouse *Differentiating OPC* gene pattern 25 projected on the mouse scRNA-seq dataset and on the human snRNA-seq dataset. **f.** Dot plot of *HIF1A* and *CTNNB1* expression in quiescent OPCs in the human cortex across aging.

**Extended Data Figure 5.**
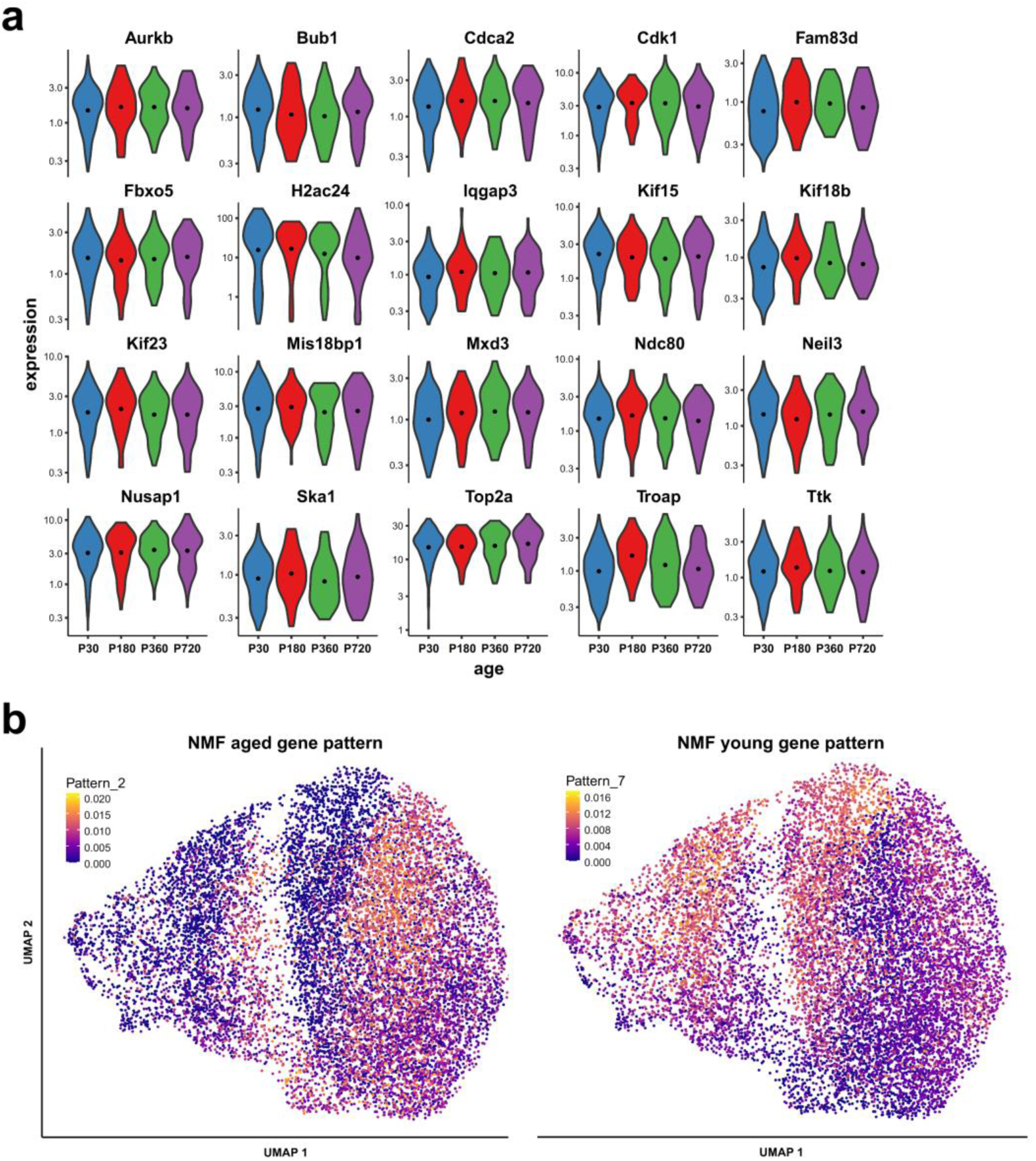
Expression changes in individual cycling genes and groups of genes in *Cycling OPC 1* and directly anteceding *Quiescent OPC*. **a.** Expression levels of known cycling genes enriched in cycling OPCs are comparable in *Cycling OPC 1* throughout aging. **b.** NMF gene patterns that are associated with either aged (P180-720) or young (P30) OPCs.

**Extended Data Figure 6.**
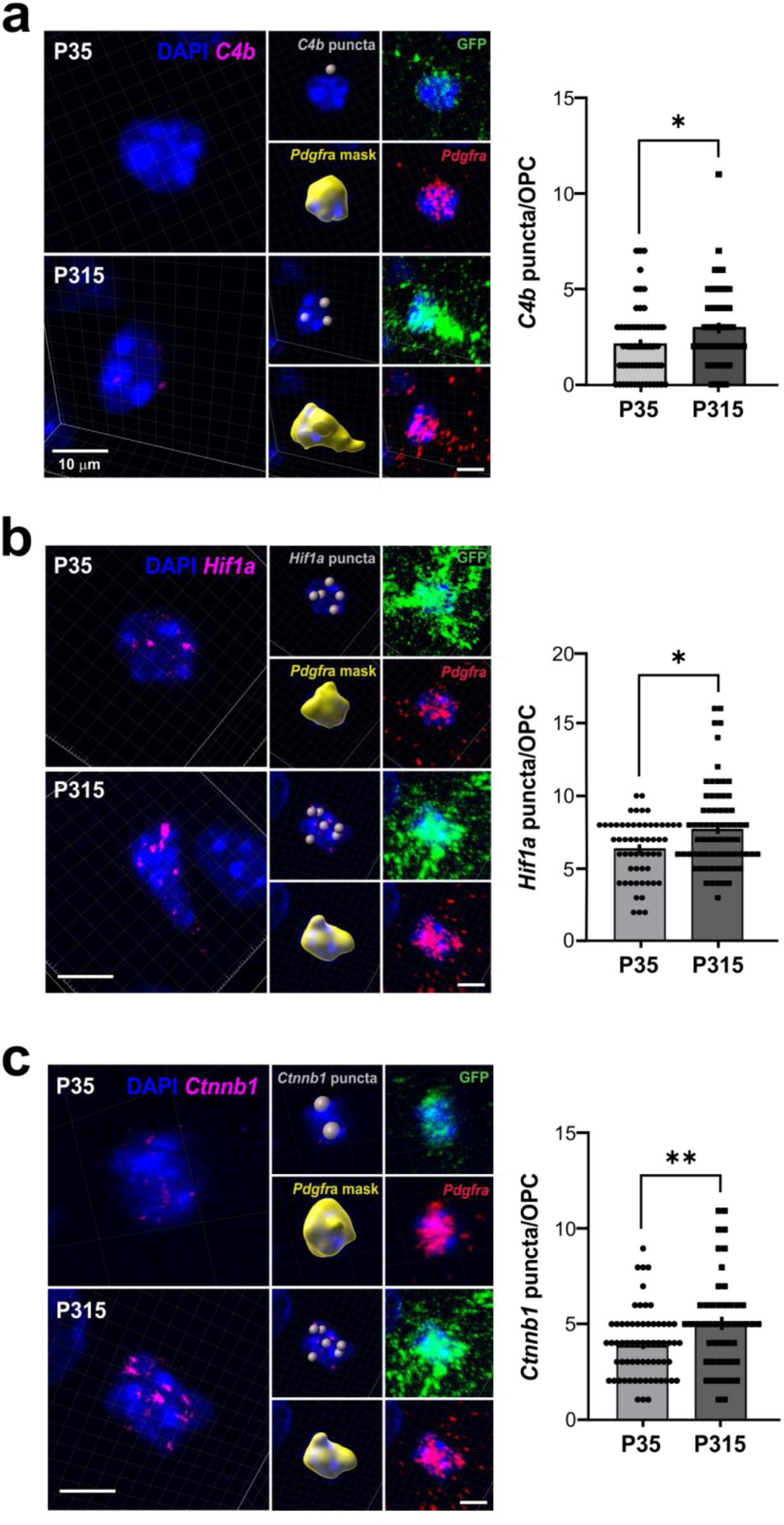
OPCs upregulate *C4b*, *Hif1a,* and *Ctnnb1* mRNA with aging. **a.** FISH with immunofluorescence staining (IF) for *Pdgfra* and *C4b* in P35 and P315 *Matn4-mEGFP* mouse cortex. OPC cell body masks were created based on EGFP fluorescence and *Pdgfra* FISH signal and used to quantify *C4b* transcript puncta/OPC (one-way ANOVA, * p-value < 0.05). **b.** FISH with IF staining for *Pdgfra* and *Hif1a* in the P35 and P315 *Matn4-mEGFP* mouse cortex. *Hif1a* transcript puncta/OPC was quantified as described above (one-way ANOVA, * p-value < 0.05). **c.** FISH with IF staining for *Pdgfra* and *Ctnnb1* in the P35 and P315 *Matn4-mEGFP* mouse cortex (one-way ANOVA, ** p-value < 0.01).

**Extended Data Figure 7.**
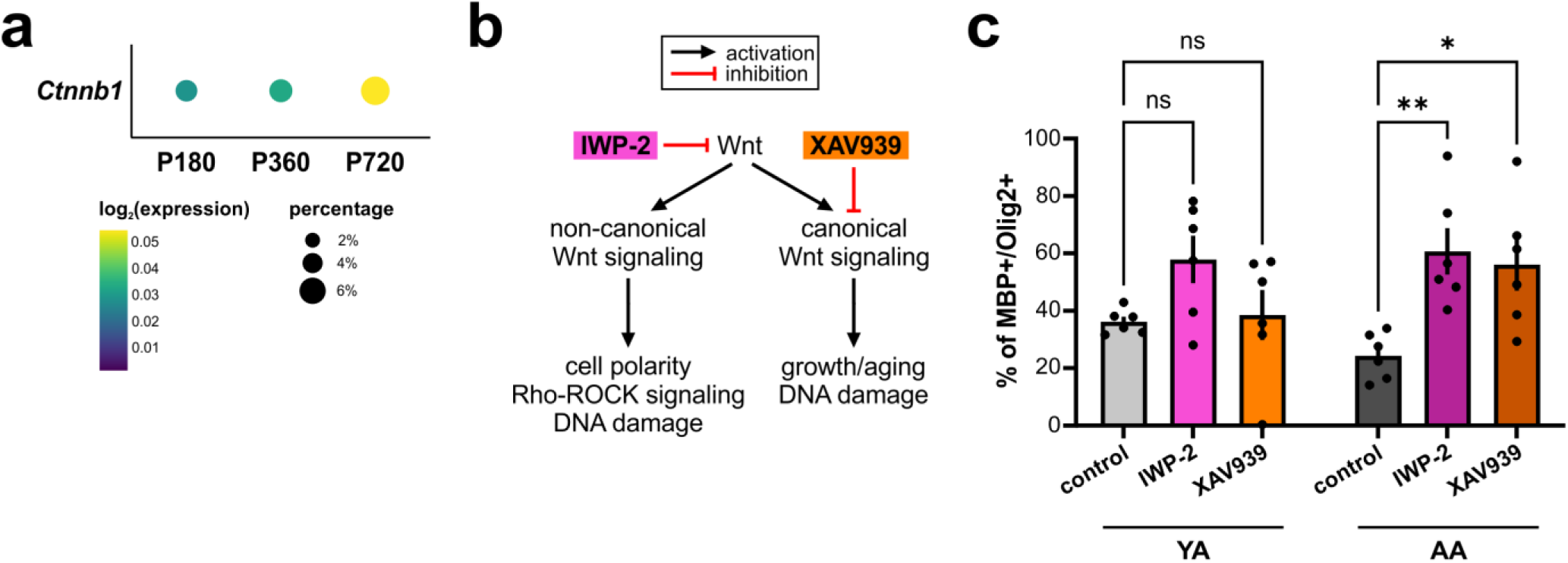
Wnt signaling pathway is activated in aged OPCs and may contribute to their decreased differentiation potential. **a.** Dot plot of *Ctnnb1* expression in *Quiescent OPC* from P180, P360, and P720 timepoints. **b.** Schematic of how two different Wnt inhibitors (IWP-2 and XAV939) differentially block Wnt signaling pathway. IWP-2 globally inhibits the Wnt pathway whereas XAV939 preferentially inhibits the canonical Wnt signaling pathway. Both non-canonical and canonical Wnt signaling pathways have been shown to regulate DNA damage response. **c.** Quantification of MBP+ differentiating OPC/Olig2+ oligodendrocyte proportions with and without Wnt inhibitor treatments in OPCs isolated from YA or AA (two-way ANOVA with Tukey’s multiple comparisons test, * p-value < 0.05, ** p-value < 0.01).

**Extended Data Figure 8.**
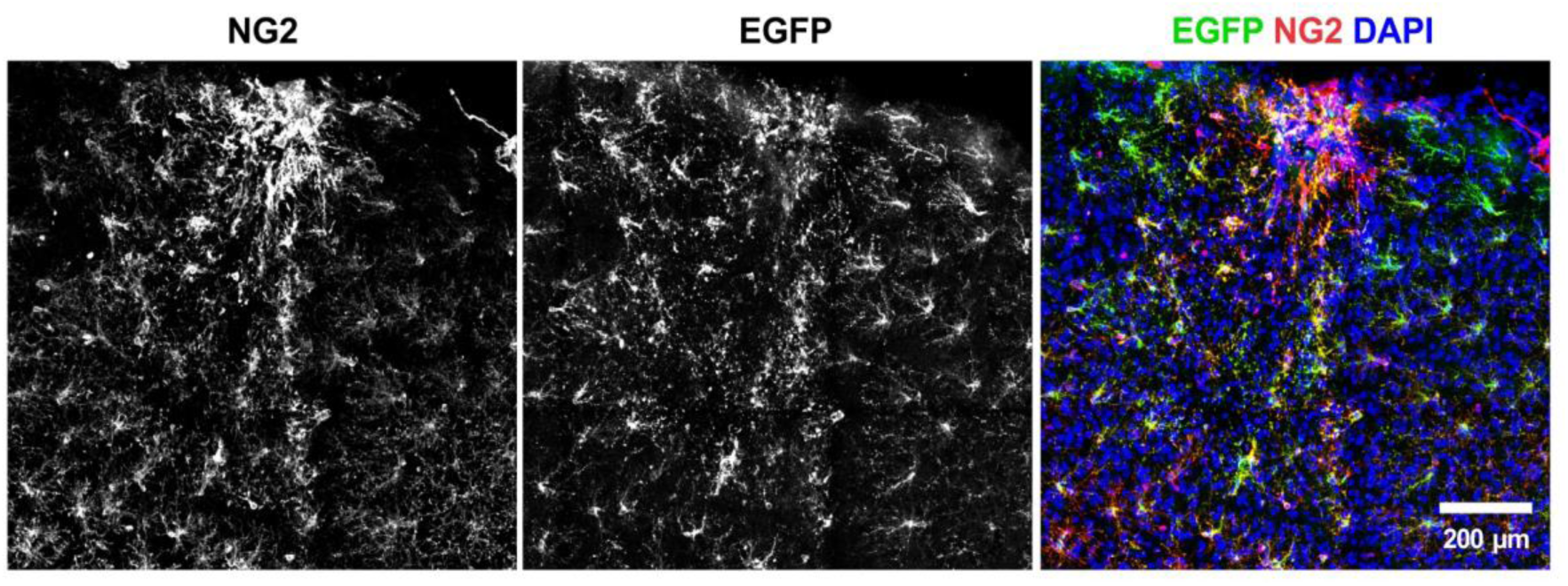
*Matn4-mEGFP* signal is restricted to OPCs even after a stab wound injury. IHC against NG2 (red) and EGFP (green) was performed on the *Matn4-mEGFP* mouse following a stab wound injury to demonstrate the utility of the mouse line in studying OPC dynamics following injury and inflammation.

**Supplementary Data 1 – Final *Matn4*-mEGFP repair template plasmid with left and right homology arms for CRISPR/Cas9.**

**Supplementary Data 2 – List of marker genes used to define OPC subtypes.**

**Supplementary Data 3 – List of genes corresponding to different gene modules associated with Quiescent, Transitioning, and Differentiating OPC clusters.**

**Supplementary Data 4 – Pseudotime differential gene expression (graph_test) of genes that encode transcription factors (TFs) along the OPC differentiation trajectory.**

**Supplementary Data 5 – Differential gene expression results of Quiescent OPCs in aging (P30 vs. P180, P360, and P720).**

**Supplementary Data 6 – Gene weights for Cycling and Differentiating OPC subtypes. Supplementary Data 7 – Quantification of *in vitro* pharmacological experiments.**

**Supplementary Video 1 – 1-hour time-lapse imaging of OPCs in the motor cortex of *Matn4*-mEGFP.**

**Supplementary Video 2 – Z-stack image of the OPCs pseudocolored in green (at Baseline of imaging) and magenta (50 minutes after Baseline).**

