## Supplementary figures and images for "Transcriptional profiles of murine oligodendrocyte precursor cells across the lifespan"

### Supplementary Data 1

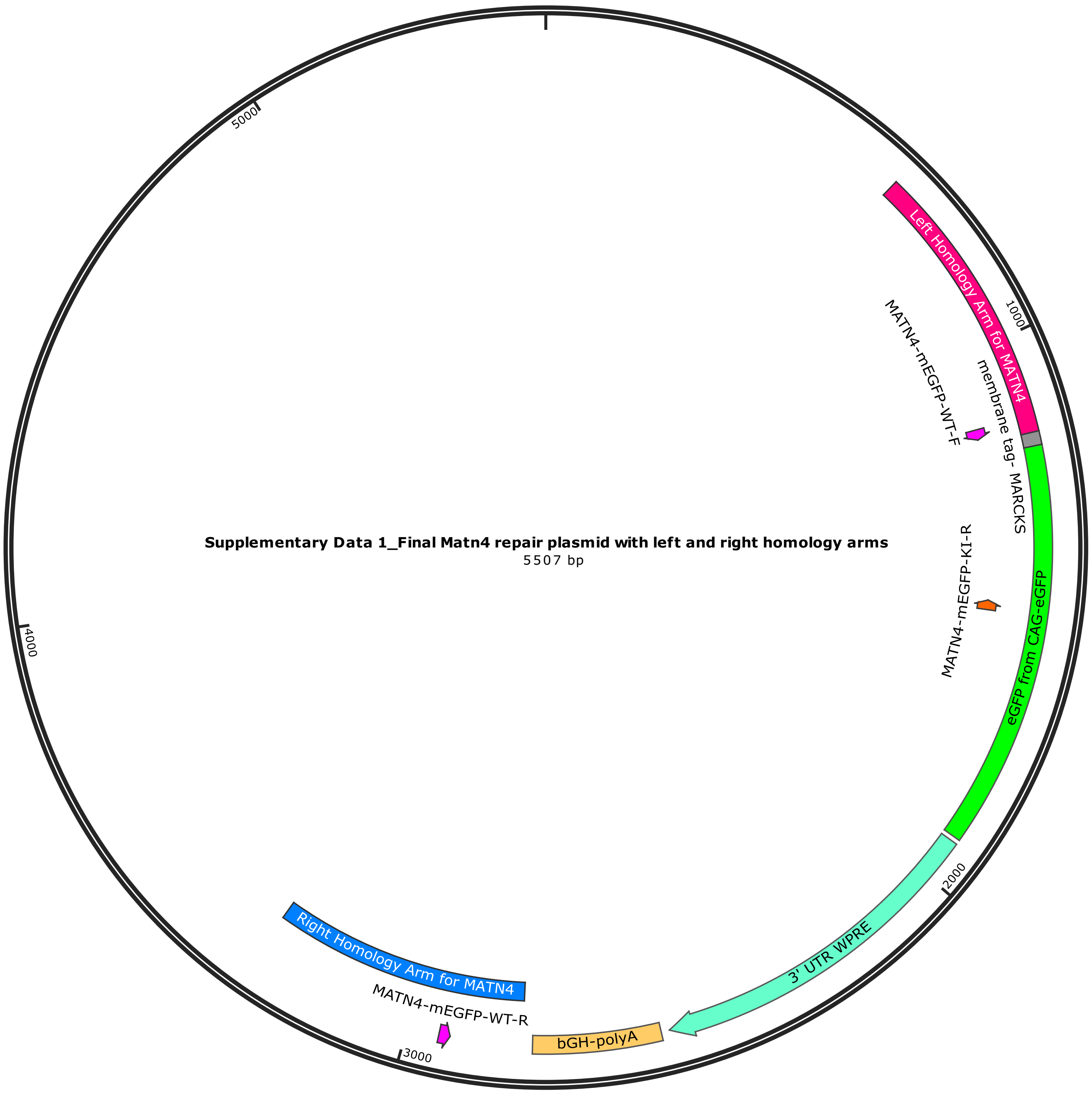

Created with SnapGene®
